# Integrative analysis of multi-omics and machine learning highlighted an m6A-related mRNA signature as a robust AAA progression predictor

**DOI:** 10.1101/2023.09.25.559437

**Authors:** Yuchen He, Jia Xing, Shiyue Wang, Han Jiang, Yu Lun, Yanshuo Han, Philipp Erhart, Dittmar Böckler, Shijie Xin, Jian Zhang

## Abstract

**Objective:** Abdominal aortic aneurysm (AAA) is a life-threatening disease in vascular surgery with significant morbidity and mortality rates upon rupture. Despite surgical interventions, effective targeted drugs for non-surgical candidates are lacking. M6A methylation, a dynamic RNA modification, has been implicated in various diseases, but its role in AAA remains poorly understood. In this study, we aimed to explore the participation of M6A in the progression of AAA progression through multi-omics and machine learning.

**Approach and Results:** we conducted methylated RNA immunoprecipitation with next-generation sequencing (MeRIP-seq) to profile the m6A methylome in AAA tissues, identifying differentially methylated genes (DMGs). Integrating multi-omics data from RNA-sequencing (RNA-seq) in GEO databases, we developed a machine learning-based AAA m6A-related mRNA signature (AMRMS) to predict AAA dilation risk. The AMRMS demonstrated robust predictive performance in distinguishing AAA patients with large AAA and small AAA. Notably, the AMRMS highlighted FKBP11 as a key gene with a significant impact on the predicted model. Subsequent single-cell RNA sequencing (ScRNA-seq) revealed the pivotal role of FKBP11-positive plasma cells in AAA progression.

**Conclusions:** Our study provides novel insights into the regulatory role of m6A modification in AAA pathogenesis, and further develop a promising AMRMS for risk evaluation in AAA patients. Furthermore, the identification of FKBP11 positive plasma cells as significant contributors to AAA progression opens new avenues for targeted therapeutic interventions.

## Introduction

Abdominal aortic aneurysm (AAA) is a prevalent degenerative vascular dilatory disease, affecting approximately 1%–2% of older individuals [1, 2]. Patients with AAA may remain asymptomatic, however hospitalized patients regularly under critical condition since AAA is progressive and its rupture can lead to catastrophic consequences, including acute abdominal pain, shock, and even death. Despite advance of open surgery and endovascuar repair, there is a lack of effective targeted drugs to prevent the progression or rupture of AAA in patients unless the diameter of the aneurysm of the patients is large enough for meeting the surgical criteria [2]. Consequently, a deeper understanding of the cellular and molecular mechanisms underlying AAA pathogenesis is urgently needed.

M6A methylation, characterized by the methylation of adenosine nucleotides in RNA molecules, has been increasingly recognized in influencing the metabolism of RNA such as RNA stability, transport, translation, and splicing [3]. By regulating RNA metabolism dynamically, m6A has been validated to promote apotosis, extracellular matrix remodeling, and inflammatory cell infiltration [4–6]. These pathological changes also involved in pathogenesis of AAA [5,7]. Actually, our team has found that m6A methylation modifications exist during the progression of AAA [8], however its role and regulation mechanism remain a mystery.

In this study, we conducted a comprehensive profiling of the m6A methylome in AAA tissue samples using Methylated RNA Immunoprecipitation with Next-Generation Sequencing (MeRIP-seq), which provided a transcriptome-wide map of m6A modifications in AAA tissues. By identifying differentially methylated genes (DMGs) and their associations with AAA pathogenesis, we aimed to disclose the underlying molecular mechanisms. Leveraging data from the AAA database in GEO, we further attempted to develop and validate a risk signature based on AAA m6A-related mRNA, enabling us to predict the risk of AAA dilation. Notably, our findings highlighted FKBP11 as the mRNA with the highest weight in the predicted model. To gain deeper insights into the cellular location and biological function of FKBP11 positive cells, we further performed single cell RNA sequense (scRNA-seq) analysis and morphological detection of tissues from AAA patients and the heathy controls. This research provides predictive models that discern m6A methylation patterns associated with disease progression, while simultaneously identifying potential therapeutic targets for managing AAA. By unraveling the intricate involvement of m6A methylation in AAA pathogenesis, our study contributes to advancing the field of vascular biology and initiates a novel treatment strategy to combat this life-threatening disease.

## Methods

### Patients and Samples

Ten patients with atherosclerotic AAA were consecutively enrolled in this study. Tissue specimens from the AAA patients and their corresponding clinical data were collected after aneurysmal open surgery repair at the First Hospital of China Medical University. Patients with Ehlers–Danlos syndrome, Marfan syndrome, other known vascular or connective tissue disorders, cancer, infection, and any other immune-related disease were excluded. Among the collected AAA tissue samples, three were used for MeRIP-seq, and the remaining seven samples were used for further scRNA-seq.

Simultaneously, seven control aortic tissue samples were obtained from organ donors serving as the normal aorta control group (three for MeRIP-seq and four for scRNA-seq). These control samples had a relatively healthy peripheral vascular system, determined by Computed Tomography Angiography (CTA), and no evidence or medical history of aneurysm or other vascular disorders. For MeRIP-seq, AAA tissue samples and matched normal tissues were promptly collected and separated into centrifuge tubes. For scRNA-seq, AAA tissue samples and control samples were stored in tissue preservation solution purchased from SeekGene BioSciences Co., Ltd., Beijing, China, and transferred under 4℃ to the SeekGene BioSciences Co. laboratory. This study was approved by the Ethics Committee of China Medical University (AF-SOP-07-1.1-01).

### RNA Extraction, MeRIP Experiment, Library Preparation, and Sequencing

MeRIP experiments and high-throughput sequencing and data analysis were conducted by Seqhealth Technology Co., Ltd. (Wuhan, China). Total RNAs were extracted from three AAA samples and three healthy control (HC) samples using TRIzol Reagent (Invitrogen, United States) following the manufacturer’s protocol. Subsequently, DNA digestion was carried out after RNA extraction using DNaseI.

RNA quality was assessed by examining A260/A280 with a Nanodrop spectrophotometer (Thermo Fisher Scientific Inc., United States), and RNA integrity was confirmed by 1.5% agarose gel electrophoresis. Qualified RNAs were quantified using Qubit3.0 with the QubitTM RNA Broad Range Assay kit (Life Technologies, United States).

For polyadenylated RNA enrichment, 50µg of total RNAs was used with VAHTS mRNA Capture Beads (Vazyme, China). RNA fragments were mainly distributed in the 100-200 nt range after the addition of 20mM ZnCl2 and incubation at 95℃ for 5-10 minutes. Then, 10% of RNA fragments were saved as “Input,” and the remaining fragments were used for m6A immunoprecipitation (IP) using a specific anti-m6A antibody (Synaptic Systems, 202203, Germany). RNA samples from both input and IP were prepared using TRIzol reagent (Invitrogen). The stranded RNA sequencing library was constructed using the KC-DigitalTM Stranded mRNA Library Prep Kit for Illumina (Seqhealth, China) following the manufacturer’s instructions. The kit employed unique molecular identifiers (UMI) of 8 random bases to label pre-amplified cDNA molecules, thereby eliminating duplication bias in PCR and sequencing steps. The library products, corresponding to 200-500 bps, were enriched, quantified, and finally sequenced on a Novaseq 6000 sequencer (Illumina) with PE150 model.

### Data Analysis for MeRIP-seq

Raw sequencing data were initially filtered by Trimmomatic (version 0.36) to discard low-quality reads and trim reads contaminated with adaptor sequences. Clean reads underwent further processing with in-house scripts to eliminate duplication bias introduced during library preparation and sequencing. Clean reads were clustered based on UMI sequences, grouping reads with the same UMI sequence together. Within each cluster, pairwise alignment was performed to identify reads with sequence identities exceeding 95%, which were then extracted into new sub-clusters. Multiple sequence alignment was conducted to obtain a consensus sequence for each sub-cluster, eliminating errors and biases introduced by PCR amplification or sequencing.

The de-duplicated consensus sequences were used for m6A site analysis, mapping to the reference genome of Homo sapiens from NCBI using STAR software (Version 2.5.3a) with default parameters. The exomePeak (Version 3.8) software was employed for peak calling, and m6A peaks were annotated using bedtools (Version 2.25.0). Differentially methylated peaks were identified using a python script with the Fisher test. Sequence motifs enriched in m6A peak regions were determined using Homer (version 4.10). Differentially methylated genes (DMGs) and specific methylated genes (SMGs) were identified using the corresponding MeRIP-seq input library data by the R package “Ballgown”. Gene Ontology (GO) analysis and Kyoto Encyclopedia of Genes and Genomes (KEGG) enrichment analysis were performed based on annotated genes using KOBAS software (version: 2.1.1) with a corrected p value cutoff of 0.05 for statistically significant enrichment.

### Public Databases and Analysis

Three mRNA sequencing datasets of humans were downloaded from the GEO database (https://www.ncbi.nlm.nih.gov/geo/), including GSE57691, GSE7084, and GSE98278. A total of 82 AAA tissue samples and 12 normal aortic tissue samples were selected and further normalized and integrated for analysis. Principal Component Analysis (PCA) was conducted using the “mixOmics” R package to detect the independence of the AAA group and the control group. The stromal score and immune score differences between the two groups were evaluated using the “Estimate” R package. Differential Expressed Genes (DEGs) between the two groups were identified with a statistical threshold of |log2Fold Change (FC)| > 1 and P < 0.05 using the “tinyarray” R package. DEGs and DMGs were then overlapped, and the intersecting genes were considered as hub genes. A heatmap diagram was constructed to display the expression of hub genes in AAA and normal aortic tissues.

Currently, twenty-one genes, including ALKBH5, FTO, HNRNPA2B1, HNRNPC, IGF2BP1, IGF2BP2, IGF2BP3, FMR1 METTL14, METTL3, RBM15, RBM15B, CBLL1, WTAP, YTHDC1, YTHDC2, YTHDF1, YTHDF2, YTHDF3, ZC3H13, and ELAVL1, are recognized as common m6A RNA methylation regulators [9]. A boxplot was generated using the “tinyarray” R package to visualize the expression of these m6A regulators in the two groups.

### Signature Generated from Machine Learning-Based Integrative Approaches

Ten machine learning algorithms, including Elastic Network (Enet), Ridge, Generalized Boosted Regression Modeling (GBM), Support Vector Machine (SVM), Random Forest (RF), Linear Discriminant Analysis (LDA), Generalized Linear Model Boosting (glmBoost), eXtreme Gradient Boosting (XGBoost), Naive Bayes, and Step Generalized Linear Model (Stepglm) were combined to identify the most robust AAA dilation signature with the highest C-index in the training dataset (GSE57691) and validating dataset (GSE98278). ROC analyses were performed for validation, and the algorithm’s prediction ability was measured using the AUC. A two-sided P < 0.05 was used to determine statistical significance. A final predictive AAA m6A-related mRNA signature (AMRMS) with the best performance was established based on the GBM algorithms and identified the 21 most valuable AMRMS.

### Cell Preparation for scRNA-seq

After harvesting, tissues were washed in ice-cold RPMI1640 and dissociated using the multi-tissue dissociation kit 2 (Miltenyi, Germany) according to the manufacturer’s instructions. DNase treatment was optional, depending on the viscosity of the homogenate. Cell count and viability were estimated using a fluorescence Cell Analyzer (Countstar Rigel S2) with AO/PI reagent after removing erythrocytes (Miltenyi) and any debris and dead cells were removed if necessary (Miltenyi). Finally, fresh cells were washed twice in RPMI1640 and resuspended at 1×10^6^ cells/ml in 1×PBS and 0.04% bovine serum albumin.

### ScRNA-seq Library Construction and Sequencing

In this study, we utilized SeekGene’s SeekOne® MM Single Cell 3’ library preparation kit to construct scRNA-seq libraries with improved efficiency and reduced duplication rate. We began by loading the appropriate number of cells into the flow channel of the SeekOne® MM chip, containing 170,000 microwells, using gravity to aid cell settling. After allowing sufficient time for settling, any remaining unsettled cells were carefully removed.

To label individual cells, we employed Cell Barcoded Magnetic Beads (CBBs), which were pipetted into the flow channel and precisely localized within the microwells using a magnetic field. Subsequently, we lysed the cells within the SeekOne® MM chip, releasing RNA that was captured by the CBBs in the same microwell. Reverse transcription was performed at 37℃ for 30 minutes to synthesize cDNA with cell barcodes. Exonuclease I treatment was used to eliminate any unused primers on the CBBs. The barcoded cDNA on the CBBs then underwent hybridization with a random primer containing a Reads 2 SeqPrimer sequence at its 5’ end, facilitating the extension and creation of the second DNA strand with cell barcodes at the 3’ end. After denaturation, the resulting second strand DNA was separated from the CBBs and purified. The purified cDNA product was amplified through PCR, and unwanted fragments were removed through a clean-up process. Full-length sequencing adapters and sample indexes were added to the cDNA through an indexed PCR. Indexed sequencing libraries were further purified using SPRI beads to remove any remaining impurities. Quantification of the libraries was performed using quantitative PCR (KAPA Biosystems KK4824). Finally, the libraries were sequenced using either the Illumina NovaSeq 6000 platform with PE150 read length or the DNBSEQ-T7 platform with PE100 read length.

### Canonical Correlation Analysis (CCA), Dimensionality Reduction, and Clustering

After quality control and filtering, library-size normalization was performed for each cell using the NormalizeData function of Seurat (version 4.0.0). Variable genes were calculated using the FindVariableGenes function. Then, all libraries were combined using FindIntegrationAnchors and IntegrateData with default parameters, and ScaleData was used to regress out the variability of the numbers of UMIs. Subsequently, RunPCA and RunTSNE were used for dimensionality reduction. FindClusters was used to cluster cells using 20 dimensions at a resolution of 0.5.

### Cell Type Identification

Cell types of different clusters were identified manually using cluster-specific marker genes from databases and literature. Each retained marker gene was expressed in a minimum of 10% of cells and at a minimum log fold change threshold of 0.25. The clustering differentially expressed genes were considered significant if the adjusted P value was less than 0.05 and the avg log2FC was more than 0.25.

### DEG Analysis for scRNA-seq

The FindMarkers function was used to identify DEGs across the conditions using the default Wilcoxon rank test. Genes were ranked by absolute log2FC, and those with P values > 0.05 (adjusted for multiple comparisons) and log2FC < 0.25 were removed.

### Enrichment Analysis for scRNA-seq

GO enrichment analysis of DEGs was implemented using the “clusterProfiler” R package. GO terms with corrected P-values less than 0.05 were considered significantly enriched by DEGs. The “clusterProfiler” R package was also used to test the statistical enrichment of DEGs in KEGG pathways.

### SCENIC Analysis and Protein-Protein Interactions (PPI) Network Construction

The Single-Cell Regulatory Network Inference and Clustering method, pySCENIC (Version 0.10.4), were used to infer transcription factor gene regulatory networks. The “pyscenic grn” function was used to generate co-expression gene regulatory networks, while AUCell analysis was performed using the “pyscenic aucell” function.

In this study, the PPI network was built using two calculation algorithms (top 5 specific regulons and their target genes) and visualized using Cytoscape software (http://cytoscape.org/).

### Pseudotime Analysis

Single-cell pseudotime trajectory analysis was performed with the “Monocle3” package to determine potential lineage differentiation. Only plasma cells in AAA tissues were analyzed. The root was determined by the expression level of FKBP11, considered a potential risk factor of AAA.

### Cell-Cell Interactions

CellPhoneDB Python package (version 2.1.7) was used to detect ligand-receptor interactions and predict communications among different cell types. To narrow down the most relevant interactions, specific interactions classified by ligand/receptor expression in more than 10% of cells within a cell type were considered. Pairwise comparisons were performed between the included cell types.

### Immunohistochemistry (IHC) Analysis

IHC analysis was performed to investigate the expression patterns of specific biomarkers within the aortic tissue samples. Representative sections of the samples from CMU Aneurysm Biobank (CMUaB), measuring 2–3 μm in thickness, were utilized for this analysis. Consecutive slides from each specimen were meticulously prepared and subjected to incubation with appropriate antibodies to precisely determine the expression of the biomarkers in individual cells within the aneurysmal lesions.

In each case, one slide was stained using an antibody to detect the specific cell type present, while a consecutive slide was stained using the antibody targeting the individual biomarker of interest. Paraffin sections were routinely stained for IHC analysis, following well-established protocols as previously described.

The primary antibodies employed in this study included an anti-J Chain antibody (dilution 1:5000; Husbio, Wuhan, China), utilized for the detection of plasma cells, and an anti-FKBP11 antibody (dilution 1:200; Absin, Shanghai, China), used to specifically target FKBP11 expression.

## Results

The flow chart of the current research was shown in **Figure 1**.

**Figure 1.**
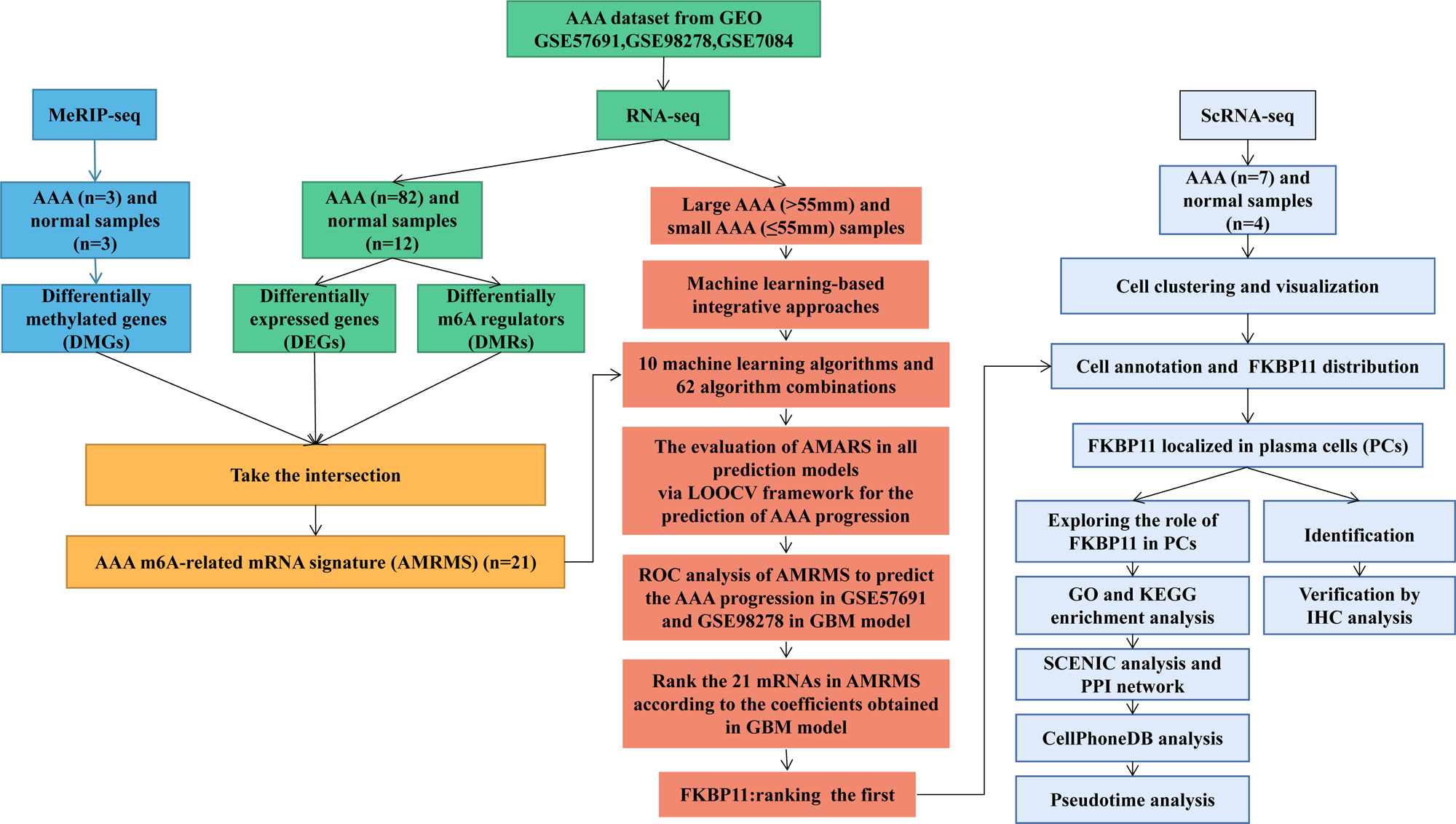
Flow chart of the research process.

### Overview of the m6A methylation map in the AAA group and the NC group

We conducted MeRIP-seq analysis on tissue samples from patients with AAA and healthy aortic tissue samples as the control group. The baseline of participants were shown in the **Table 1**. Our results revealed a total of 6988 m6A peaks in the AAA group and 7256 m6A peaks in the NC group (**Figure 2A**). Among these peaks, 5947 m6A peaks were detected in both groups, while 1309 and 1041 m6A peaks were specifically detected in the AAA and NC groups, respectively, as illustrated in the Venn diagram (**Figure 2B**).

**Figure 2.**
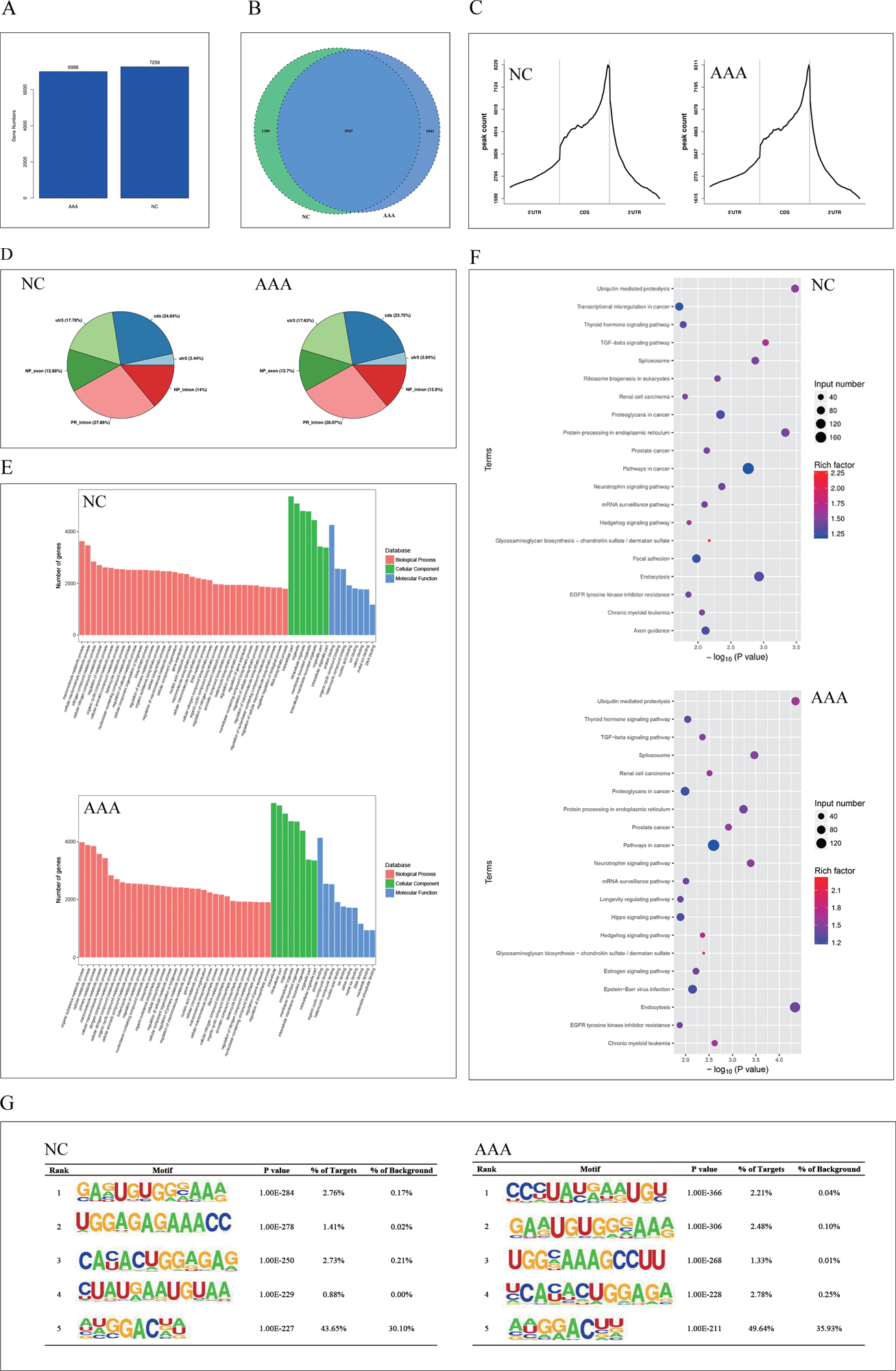
Transcriptome-wide MeRIP-seq and analysis of m6A peaks. (**A**) Overlap of m6A peaks in the AAA group and the normal control (NC) group. (**B**) Numbers of AAA-unique, NC-unique, and common m6A peaks are shown as Venn diagram as well as m6A peaks representing genes in the two groups. (**C**) and (**D**) Proportion of m6A peaks distributed in the indicated regions in the NC group and the AAA. (**E**) Major gene ontology terms significantly enriched in the two groups. (**F**) Bubble plots of major KEGG terms significantly enriched in the two groups. (**G**) Top 5 m6A modification motifs enriched from all identified m 6 A peaks in the two groups.

**Table 1.**
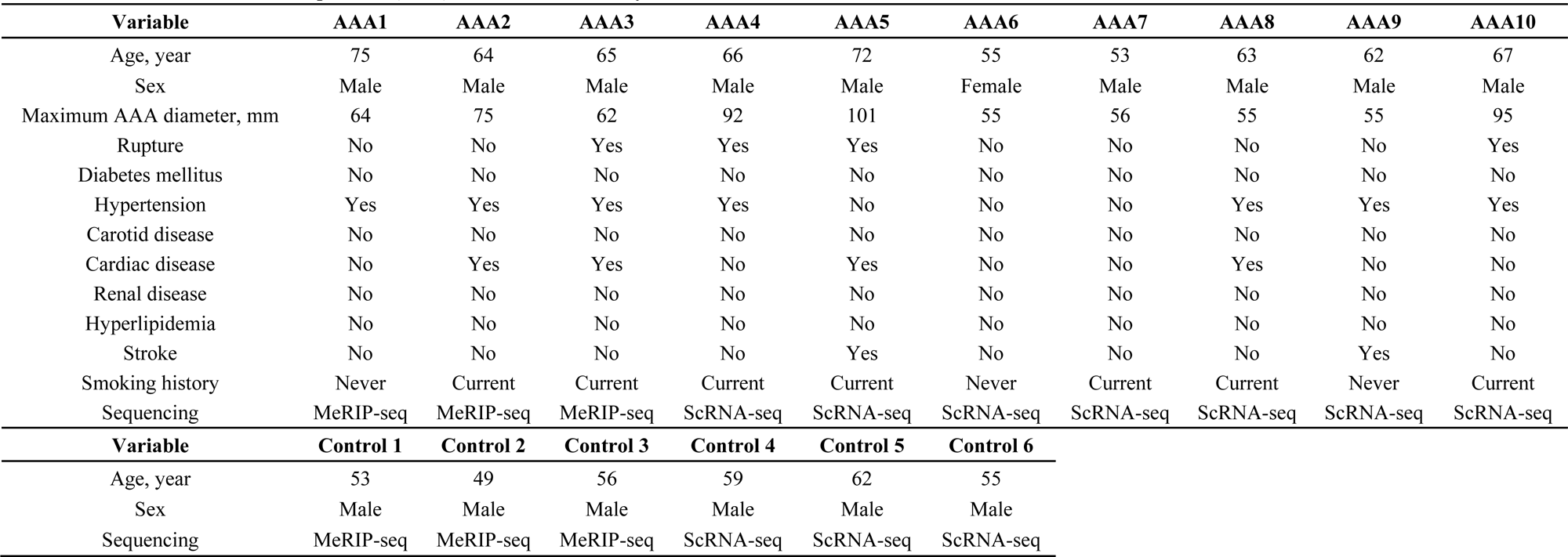
The baseline data of AAA patients (n=10) included in this study.

Furthermore, we explored the distribution of m6A peaks in both NC and AAA tissues. The m6A peaks in both tissues were predominantly enriched in the coding sequence (CDS) near the 3’ untranslated region (3’UTR) of the stop codon (**Figure 2C** and **2D**). Notably, the pattern of m6A peaks in CDS (23.75% vs. 24.04%), 3’UTR (17.63% vs. 17.73%), and 5’ untranslated region (5’UTR) (3.94% vs. 3.44%) was similar between AAA tissues and healthy aortic tissues.

To gain insights into the biological functions of m6A modification, we performed GO and KEGG pathway analyses for m6A methylated genes in healthy aortic and AAA tissues. The GO analysis classified genes into three functional groups: biological process (BP), cellular component (CC), and molecular function (MF) (**Figure 2E**). Additionally, our KEGG pathway analysis demonstrated significant associations between m6A methylated genes and various signal pathways in both groups (**Figure 2F**).

Further analysis revealed the top five m6A motifs enriched from altered m6A peaks in both groups (**Figure 2G**). These motifs hold potential significance for understanding the regulatory mechanisms and functions of m6A methylation in the context of AAA and healthy aortic tissues.

### Differentially Methylated mRNAs Involved in Key Signaling Pathways

Comparing the NC group to the AAA group, we observed 805 significantly up-regulated m6A peaks, corresponding to transcripts of 750 genes, and 559 significantly down-regulated m6A peaks, representing transcripts of 529 genes (| log2FC | > 1 and p < 0.05). The top 20 altered up-regulated m6A peaks are presented in **Table 2**, while the top 20 altered down-regulated m6A peaks are listed in **Table 3**.

**Table 2.** Top 20 up-regulated genes in the AAA group compared with the NC group.

| Gene | Chromosome | Peak Start | Peak End | logFC | P Value | Annotation |
| --- | --- | --- | --- | --- | --- | --- |
| RP11-458D21.1 | chr1 | 146058976 | 146059187 | 8.549092311 | 3.96E-258 | Exon |
| AC006460.2 | chr2 | 190699764 | 190701481 | 8.331286675 | 9.26E-100 | Exon |
| ANKRD35 | chr1 | 145872259 | 145872349 | 6.572519874 | 3.46E-100 | CDS |
| RAB6B | chr3 | 133826000 | 133826120 | 6.103135181 | 2.35E-25 | 3'UTR |
| RP11-190A12.9 | chr1 | 159910150 | 159910267 | 6.091453376 | 1.31E-86 | Exon |
| ANKRD35 | chr | 145872617 | 145873155 | 5.252018306 | 2.52E-49 | CDS |
| PEX11B | chr1 | 145912398 | 145912549 | 4.918719817 | 2.67E-50 | CDS |
| CCNT2 | chr2 | 134918621 | 134918710 | 4.801015567 | 1.35E-17 | 5'UTR |
| SLC16A9 | chr10 | 59654199 | 59654409 | 4.529542873 | 1.71E-68 | CDS |
| RP11-956E11.1 | chr12 | 69459578 | 69460640 | 4.509641267 | 7.00E-33 | Exon |
| B9D1 | chr17 | 19377782 | 19377902 | 4.48020483 | 1.39E-17 | Exon |
| SH3BP5L | chr1 | 248825914 | 248826005 | 4.425779076 | 4.29E-07 | 5'UTR |
| IRS2 | chr13 | 109752906 | 109753057 | 4.382558547 | 2.26E-07 | 3'UTR |
| NRG1 | chr8 | 32764036 | 32764247 | 4.376053202 | 1.56E-15 | CDS |
| CLN5 | chr13 | 76991645 | 76991855 | 4.288490223 | 3.39E-08 | 5'UTR |
| PEG10 | chr7 | 94663468 | 94663528 | 4.231928235 | 4.23E-12 | CDS |
| SP4 | chr7 | 21429813 | 21429934 | 4.116758465 | 1.78E-12 | CDS |
| ABCC10 | chr6 | 43432629 | 43432690 | 4.103135181 | 1.77E-35 | CDS |
| OR1K1 | chr9 | 122800508 | 122800568 | 4.090045181 | 1.43E-32 | CDS |
| SYNGAP1 | chr6 | 33456473 | 33456623 | 4.06851564 | 3.63E-05 | 3'UTR |

**Table 3.** Top 20 down-regulated genes in the AAA group compared with the NC group.

| Gene | Chromosome | Peak Start | Peak End | logFC | P Value | Annotation |
| --- | --- | --- | --- | --- | --- | --- |
| AGTR1 | chr3 | 148742109 | 148742348 | -6.336600727 | 2.23E-157 | 3'UTR |
| C1GALT1C1L | chr2 | 43675589 | 43675824 | -5.829866907 | 1.74E-27 | CDS |
| LINC01099 | chr4 | 177907817 | 177907936 | -5.82891394 | 7.46E-35 | Exon |
| APOB | chr2 | 21012100 | 21012221 | -5.67323491 | 4.17E-61 | CDS |
| PLCXD3 | chr5 | 41382142 | 41382382 | -5.420916738 | 1.69E-29 | CDS |
| KCNK17 | chr6 | 39299326 | 39299386 | -5.141089108 | 1.07E-39 | 3'UTR |
| SACS-AS1 | chr13 | 23428362 | 23428422 | -4.892928011 | 2.22E-32 | Exon |
| MAGEC3 | chrX | 141897265 | 141897326 | -4.672031829 | 1.13E-07 | CDS |
| COLEC10 | chr8 | 119105934 | 119106141 | -4.590041585 | 2.21E-13 | CDS |
| AC011625.1 | chr7 | 132367800 | 132367859 | -4.583511243 | 2.66E-15 | Exon |
| KCTD16 | chr5 | 144207227 | 144207378 | -4.471213059 | 7.98E-23 | CDS |
| KCNJ3 | chr2 | 154854794 | 154855273 | -4.460750379 | 4.28E-06 | CDS |
| TENM1 | chrX | 124380632 | 124380993 | -4.394137407 | 7.90E-12 | CDS |
| CDH8 | chr16 | 62021340 | 62021580 | -4.233979854 | 2.09E-07 | CDS |
| RP1-95L4.4 | chr6 | 143343173 | 143343383 | -4.213221928 | 4.34E-24 | Exon |
| LLNLF-187D8.1 | chr19 | 46428291 | 46428412 | -4.18325756 | 3.30E-03 | CDS |
| CD200 | chr3 | 112332233 | 112332410 | -4.027465659 | 1.51E-06 | 5'UTR |
| ADRA1D | chr20 | 4221409 | 4221560 | -4.014672632 | 4.27E-05 | 3'UTR |
| RP11-768F21.1 | chr12 | 119594280 | 119594340 | -3.992971755 | 3.74E-12 | Exon |
| FZD3 | chr8 | 28502835 | 28503076 | -3.988564653 | 3.57E-09 | CDS |

To gain insights into the functional implications of these differentially methylated mRNAs, we conducted comprehensive GO analysis and KEGG pathway analysis for genes with up-regulated m6A peaks (**Figure 3A**). The GO analysis revealed their enrichment in essential biological processes, including positive regulation of mitophagy (GO term: BP), autophagosome (GO term: CC), and DNA binding (GO term: MF). Moreover, the KEGG analysis unveiled significant enrichment in several signaling pathways, such as the Notch signaling pathway, ABC transporters, Mucin type O-Glycan biosynthesis, and RNA degradation. Similarly, we performed GO and KEGG pathway analyses for genes with down-regulated m6A peaks (**Figure 3B**). These genes were found to be mainly enriched in positive regulation of mitophagy (GO term: BP), divalent inorganic cation homeostasis (GO term: CC), and Smooth muscle contraction (GO term: MF), which aligns with the findings in the up-regulated m6A peak group. Additionally, KEGG analysis revealed significant enrichment in signal pathways like the Ubiquitin-mediated proteolysis, Regulation of actin cytoskeleton, and Calcium signaling pathway.

**Figure 3.**
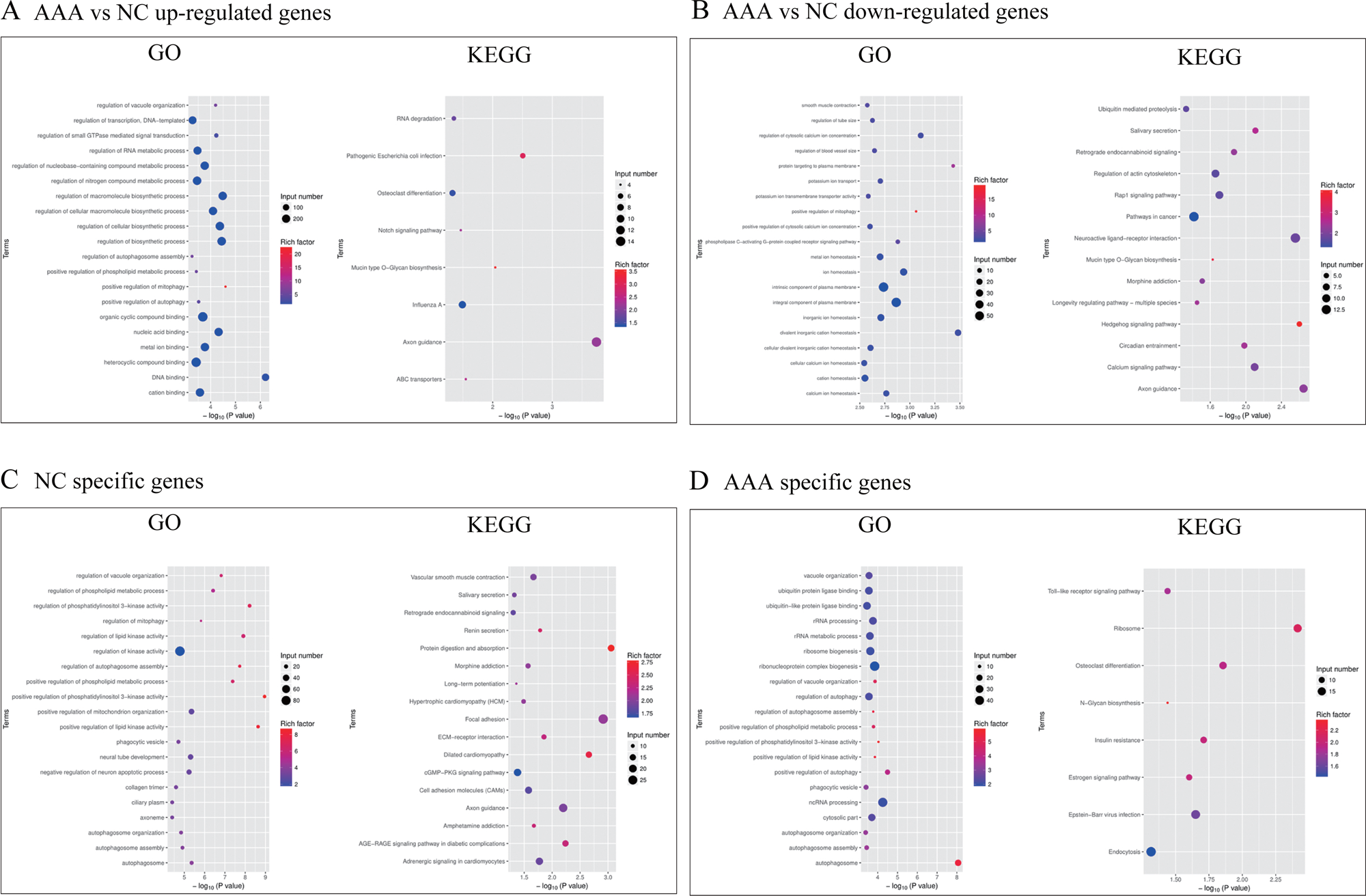
Systematic functional analysis of genes with differentially m6A modification. The Bubble plots of GO analysis and KEGG analysis for up-regulated methylated genes and down-regulated methylated genes in AAA shown in (**A**) and (**B**). Major GO enrichments and KEGG enrichments of NC specific methylated genes (**C**) and AAA specific methylated genes (**D**).

Next, we conducted GO and KEGG pathway analyses for genes with specific m6A peaks in AAA and healthy aorta (**Figure 3C** and **3D**). The AAA-specific m6A methylated genes were enriched in several biological processes, such as positive regulation of autophagy (GO term: BP), autophagosome (GO term: CC), and ubiquitin protein ligase binding (GO term: MF). Furthermore, KEGG analysis revealed significant enrichment in signal pathways like the Toll-like receptor signaling pathway, Osteoclast differentiation, Notch signaling pathway, N-Glycan biosynthesis, and Ribosome pathway. Additionally, we identified NC-specific m6A methylated genes, which were enriched in several biological processes, such as positive regulation of phosphatidylinositol 3-kinase activity (GO term: BP), autophagosome (GO term: CC), and regulation of lipid kinase activity (GO term: MF). KEGG analysis also revealed significant enrichment in signal pathways like Vascular smooth muscle contraction, Focal adhesion, and ECM-receptor interaction.

### Identification of hub DMGs and Key m6A Regulators in AAA through Conjoint Analysis of MeRIP-seq and RNA-seq Data

We performed a comprehensive analysis by integrating MeRIP-seq and RNA-seq data to identify hub DMGs in AAA. PCA clearly distinguished mRNA profiles obtained from AAA samples (n=82) and normal aortic tissue samples as controls (n=12) in the GSE57691, GSE7084, and GSE98278 datasets. To assess the differences between the two groups, we evaluated the stromal score and immune score. The results indicated that the AAA group exhibited significantly higher stromal and immune scores compared to the control group (**Figure 4A** and **4B**).

**Figure 4.**
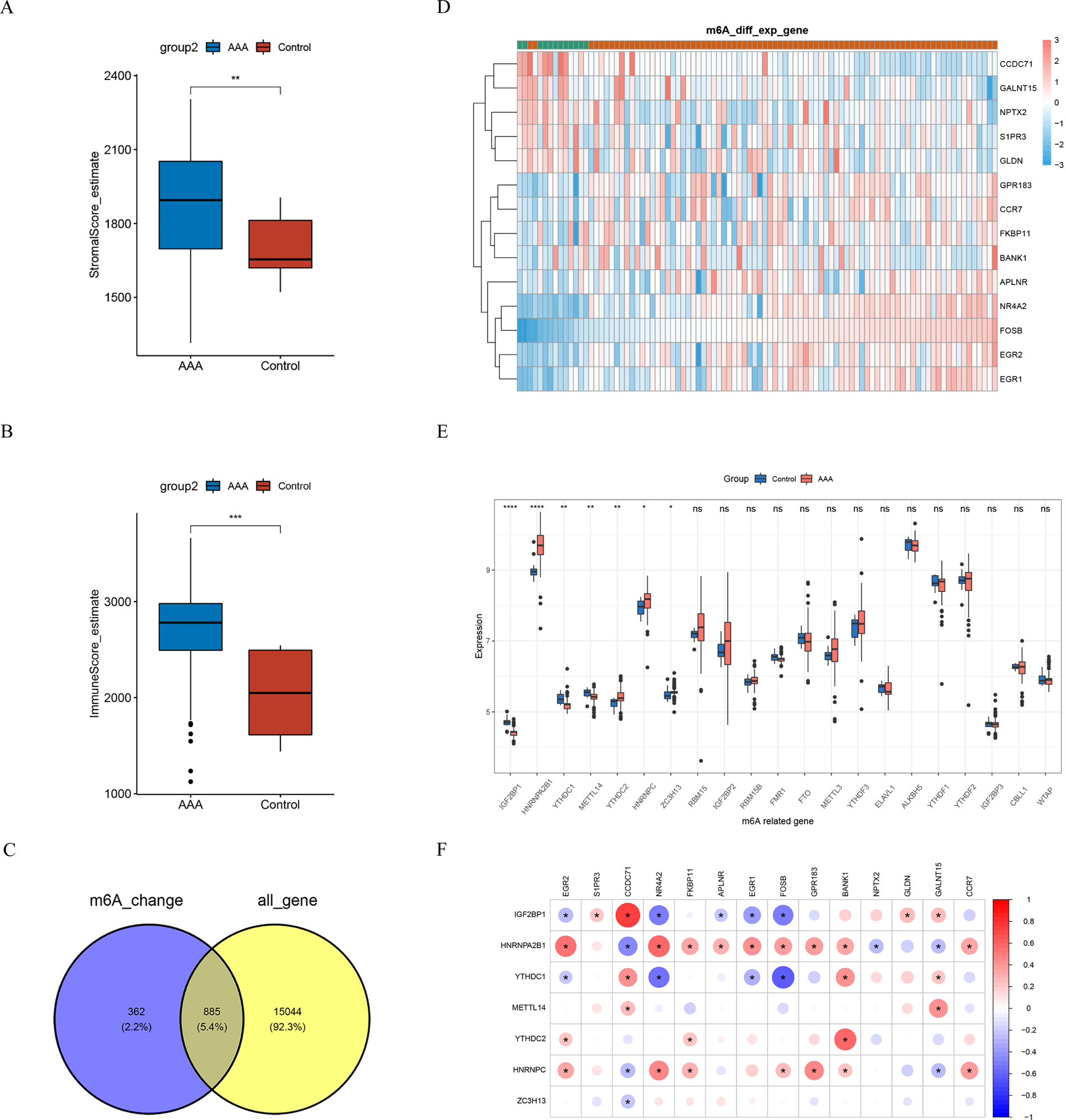
The conjoint analysis of MeRIP-seq and RNA-seq from GEO database to explore m6A expression pattern in AAA tissue samples. Comparison of stromal score (**A**) and immune score (**B**) in the AAA group and the control group shown in the histograms. (**C**) Venn diagram of DMGs from MeRIP-seq and all genes from RNA-seq. (**D**) The expressions 14of hub DMGs in 82 AAA samples and 12 control samples were shown in the heatmap. (**E**) The differential analysis of 21 m6A modulators between the AAA group and the control group was shown in the boxplots. (**F**) The correlation among hub DMGs and DMRs were shown through correlation heatmap plot.

Next, we obtained a total 885 differentially expressed DMG through overlapping all genes obtained from RNA-seq data with the DMGs identified from MeRIP-seq data (| log2FC | > 0.585 and p < 0.05). (**Figure 4C**). Then we narrowed the range of DEGs by (| log2FC | > 1 and p < 0.05). We found 14 genes that overlapped between the two datasets, which we considered as the hub DMGs (**Figure 4D** and **Table 4**). Among these hub genes, 9 genes, namely NR4A2, FKBP11, APLNR, EGR2, EGR1, FOSB, GPR183, BANK1, and CCR7, were up-regulated in the AAA group compared to the controls. On the other hand, 5 genes, including S1PR3, CCDC71, NPTX2, GLDN, and GALNT15, were down-regulated in the AAA group.

**Table 4.** Hub differentially methylated genes in AAA from the integration of GSE57691, GSE7084, and GSE98278.

| Gene | Entrez ID | Log2FC | Ave Expr | P Value | Adj P Value | Change |
| --- | --- | --- | --- | --- | --- | --- |
| FOSB | 2354 | 3.087 | 11.371 | 1.034E-12 | 4.747E-10 | up |
| NR4A2 | 4929 | 1.810 | 9.525 | 1.410E-06 | 5.325E-05 | up |
| EGR2 | 1959 | 1.740 | 8.382 | 2.312E-06 | 7.890E-05 | up |
| CCR7 | 1236 | 1.708 | 6.920 | 1.208E-03 | 1.034E-02 | up |
| EGR1 | 1958 | 1.262 | 11.512 | 2.730E-06 | 9.038E-05 | up |
| GPR183 | 1880 | 1.208 | 8.931 | 3.414E-03 | 2.222E-02 | up |
| BANK1 | 55024 | 1.102 | 5.766 | 7.335E-04 | 7.154E-03 | up |
| FKBP11 | 51303 | 1.077 | 8.925 | 1.498E-03 | 1.213E-02 | up |
| APLNR | 187 | 1.058 | 7.632 | 1.667E-04 | 2.258E-03 | up |
| CCDC71 | 64925 | -1.115 | 4.909 | 5.570E-23 | 9.204E-19 | down |
| S1PR3 | 1903 | -1.279 | 7.328 | 1.762E-04 | 2.359E-03 | down |
| GLDN | 342035 | -1.446 | 6.657 | 8.482E-04 | 7.999E-03 | down |
| GALNT15 | 117248 | -1.642 | 6.201 | 5.382E-11 | 1.202E-08 | down |
| NPTX2 | 4885 | -1.951 | 7.003 | 1.723E-06 | 6.254E-05 | down |

Note, Ave Expr, Average expression; Adj P Value, Adjustment P Value

The expression patterns of m6A regulators were investigated based on the datasets GSE57691, GSE7084, and GSE98278, as depicted in **Figure 4E**. Our analysis revealed seven differentially expressed m6A regulators (IGF2BP1, HNRNPA2B1, YTHDC1, METTL14, YTHDC2, HNRNPC, and ZC3H13) between AAA and control tissues, which we classified as differentiated m6A regulators (DMRs). Specifically, HNRNPA2B1, YTHDC2, and HNRNPC exhibited higher expression levels in AAA tissues, whereas other regulators were down-regulated in AAA tissues compared to the control aortic tissues.

To gain further insights into the regulatory interactions between significant m6A regulators and hub DMGs, we conducted correlation analyses and presented the results in **Figure 4F**. The findings shed light on the potential regulatory associations between these m6A regulators and the identified hub genes, which play pivotal roles in AAA development and progression.

### Integrative AAA m6A-Related mRNA Signature (AMRMS) comprising hub DMGs and Key m6A Regulators Predicts Large AAA Cases with High Accuracy

To develop a robust and reliable signature for AAA prediction, we employed a machine learning-based integrative approach, utilizing 14 hub DMGs and 7 DMRs. This integrative signature was referred to as AAA m6A-Related mRNA Signature.

In our analysis, we utilized two independent datasets, GSE57691 and GSE98278, comprising 29 large AAAs (lAAAs) and 20 small AAAs (sAAA), and 16 large AAAs and 15 small AAAs, respectively. All AAA samples were categorized into high-risk and low-risk groups based on their sizes in GSE57691 and GSE98278 datasets.

To evaluate the predictive performance of AMRMS specifically for large AAA, we implemented a Leave-One-Out Cross-Validation (LOOCV) framework to train and test the 113 different prediction models. Subsequently, the performance of each model was quantified using the C-index across both datasets, GSE57691 and GSE98278. (**Figure 5A** and **5B**)

**Figure 5.**
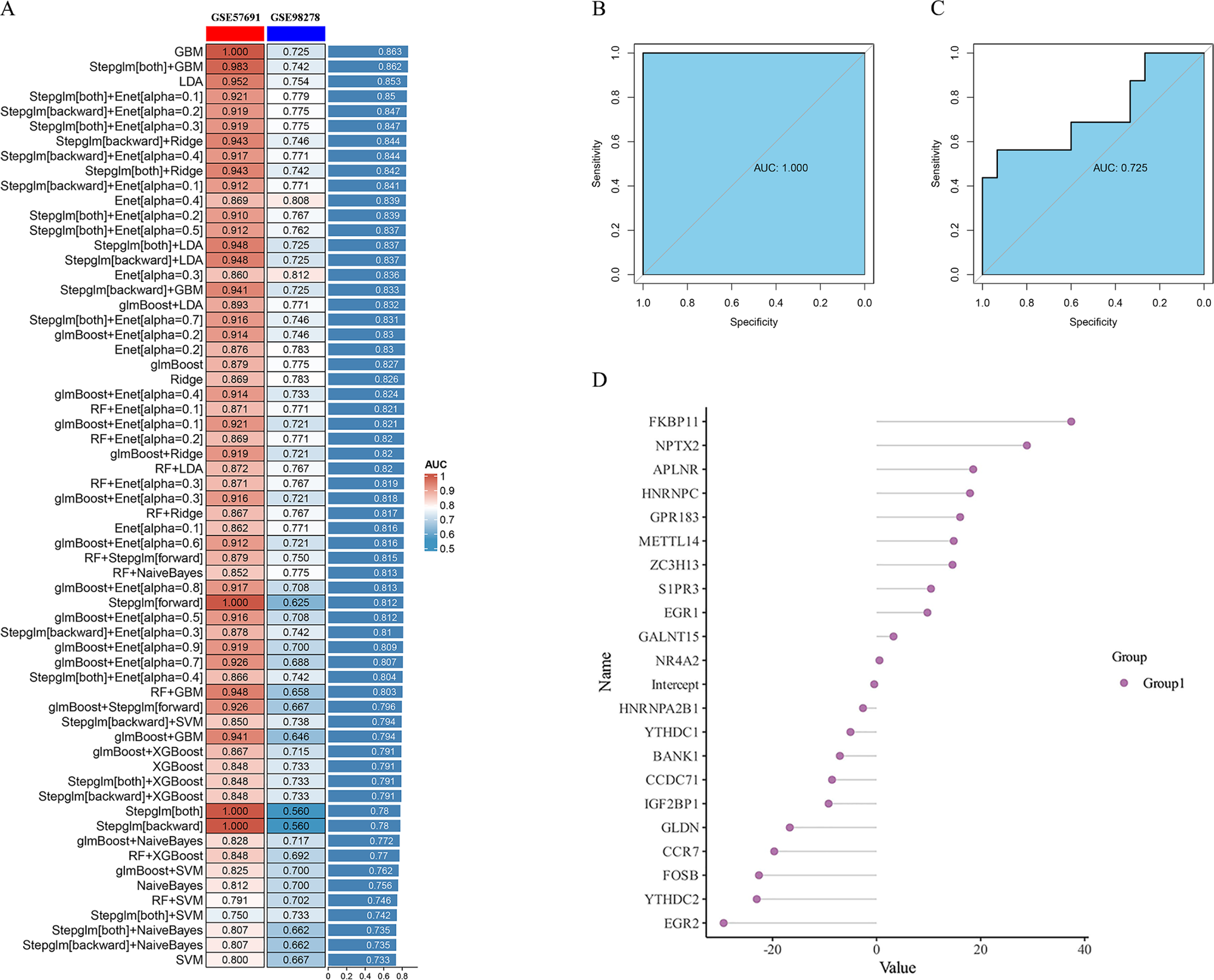
The consensus AMRMS was developed and validated to predict the risk of large AAA via the machine learning-based integrative procedure. **(A)** A total of 62 kinds of prediction models via LOOCV framework and further calculated the C-index of each model across all validation datasets. ROC curves of AMRMS to predict the risk of large AAA in GSE57691 (**B**) and GSE98278 (**C**). (**D**) Coefficients of 21 mRNAs of AMRMS finally obtained in GBM model.

Notably, the GBM model, incorporating the AMRMS gene set, outperformed other models in accurately predicting large AAA cases. Additionally, Receiver Operating Characteristic (ROC) analysis demonstrated that AMRMS effectively distinguished between large AAA and small AAA samples, yielding Area Under the Curve (AUC) values of 1.00 and 0.725 in GSE57691 and GSE98278, respectively.

To calculate a risk score for each sample, we employed the expression levels of 21 mRNAs from the AMRMS signature, weighted by their respective regression coefficients in the GBM model. Among these 21 mRNAs, FKBP11 exhibited the most positive correlation with large AAA, suggesting its potential significance in the pathogenesis of this vascular disorder.

### The expression of FKBP11 was up-regulated in large AAAs compared with that in small AAAs based on the differential expression analysis on mRNAs from AMRMS

We further compared the expressions of mRNAs from AMRMS between the lAAAgroup and the sAAA group in the training dataset (GSE57691) and the testing dataset (GSE98278). Notably, in GSE57691, we found that FKBP11 and APLNR were significantly up-regulated in lAAA (P=0.0018, and P=0.0094, respective), while IGF2BP1 was significantly down-regulated in lAAA in contrast to the sAAA group (P=0.041). (**Figure 6A**) In addition, in GSE98278, only FKBP11 was significantly up-regulated in the lAAA group in comparison to the sAAA group. Our result indicated a critical role of FKBP11 in the progression of AAA.

**Figure 6.**
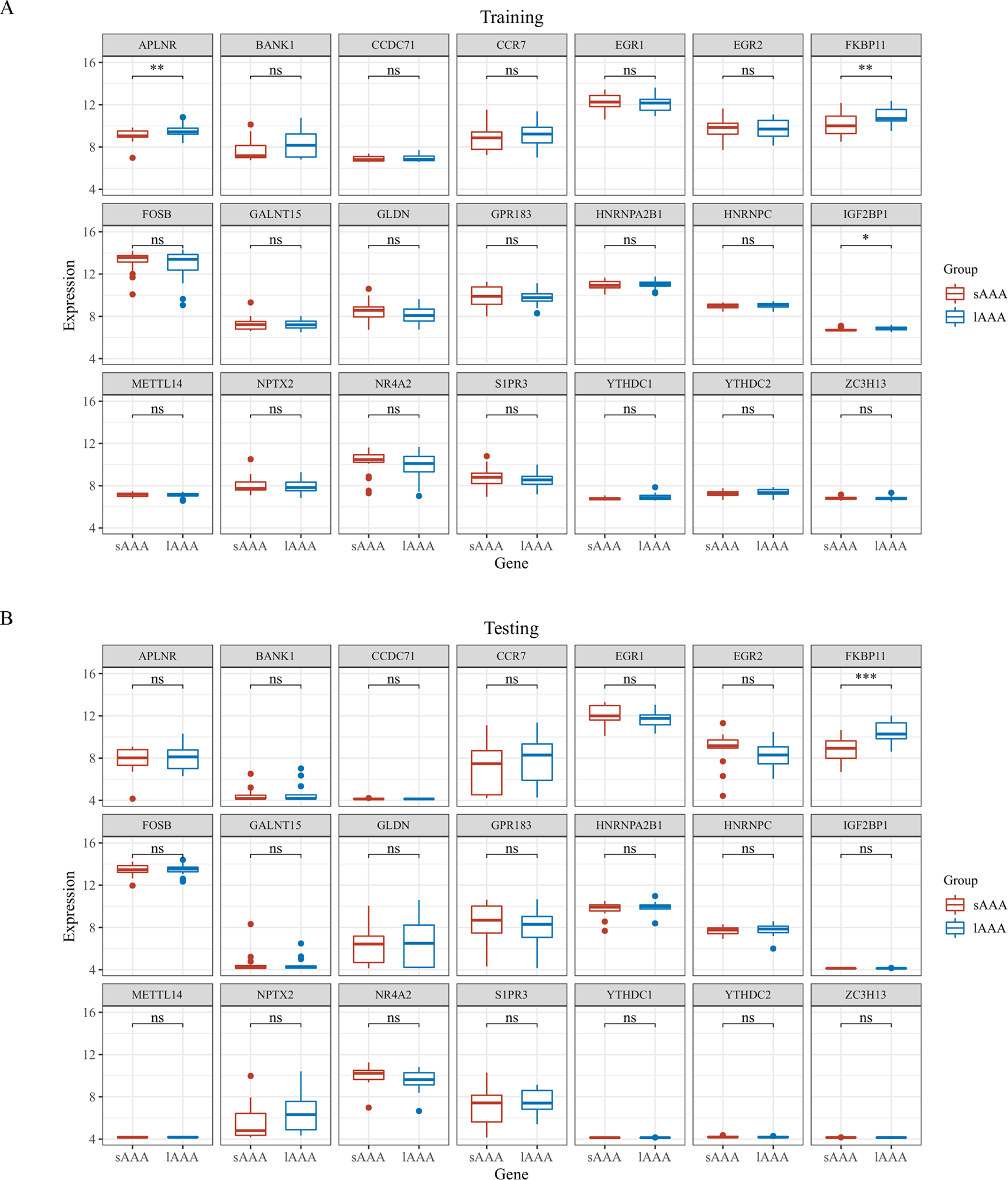
The expression of mRNAs in AMRMS in lAAA and sAAA tissue samples. (**A**) The comparison of the expression of 21 mRNAs bewteen the lAAA group and sAAA group in GSE57691 as shown in the boxplots. (**B**) The comparison of the expression of 21 mRNAs bewteen the lAAA group and sAAA group in GSE98278 as shown in the boxplots. *, P<0.05; **, P<0.01; and ***, P<0.001.

### FKBP11 is highly expressed in plasma cells residing in AAA samples based on scRNA-seq and IHC staining

Baseline clinical data of the 10 participants are presented in **Table 1**. A total of 68,326 cells were successfully recovered after applying quality control filters, comprising 37,743 cells from the AAA group and 30,583 cells from the healthy control group.

Cell populations were identified based on gene expression patterns of established canonical markers, as shown in **Table 5**. Ten distinct cell populations were manually annotated and visualized using t-Distributed Stochastic Neighbor Embedding (t-SNE) plot, with each cell type exhibiting a unique gene expression profile (**Figure 7A-C**). The top 5 enriched genes for each cell population in both AAA and control groups were displayed in a heatmap (**Figure 7C**). Major cell populations included mononuclear phagocytes (MPs), endothelial cells (ECs), fibroblasts, smooth muscle cells (SMCs), myofibroblasts. Among the non-immune cell populations, the percentage of fibroblasts (13.9% vs 5.1% in the healthy control group) and ECs (11.0% vs 5.7% in the healthy control group) were higher in AAA samples, whereas the percentage of SMCs decreased in AAA compared to the healthy control group.

**Figure 7.**
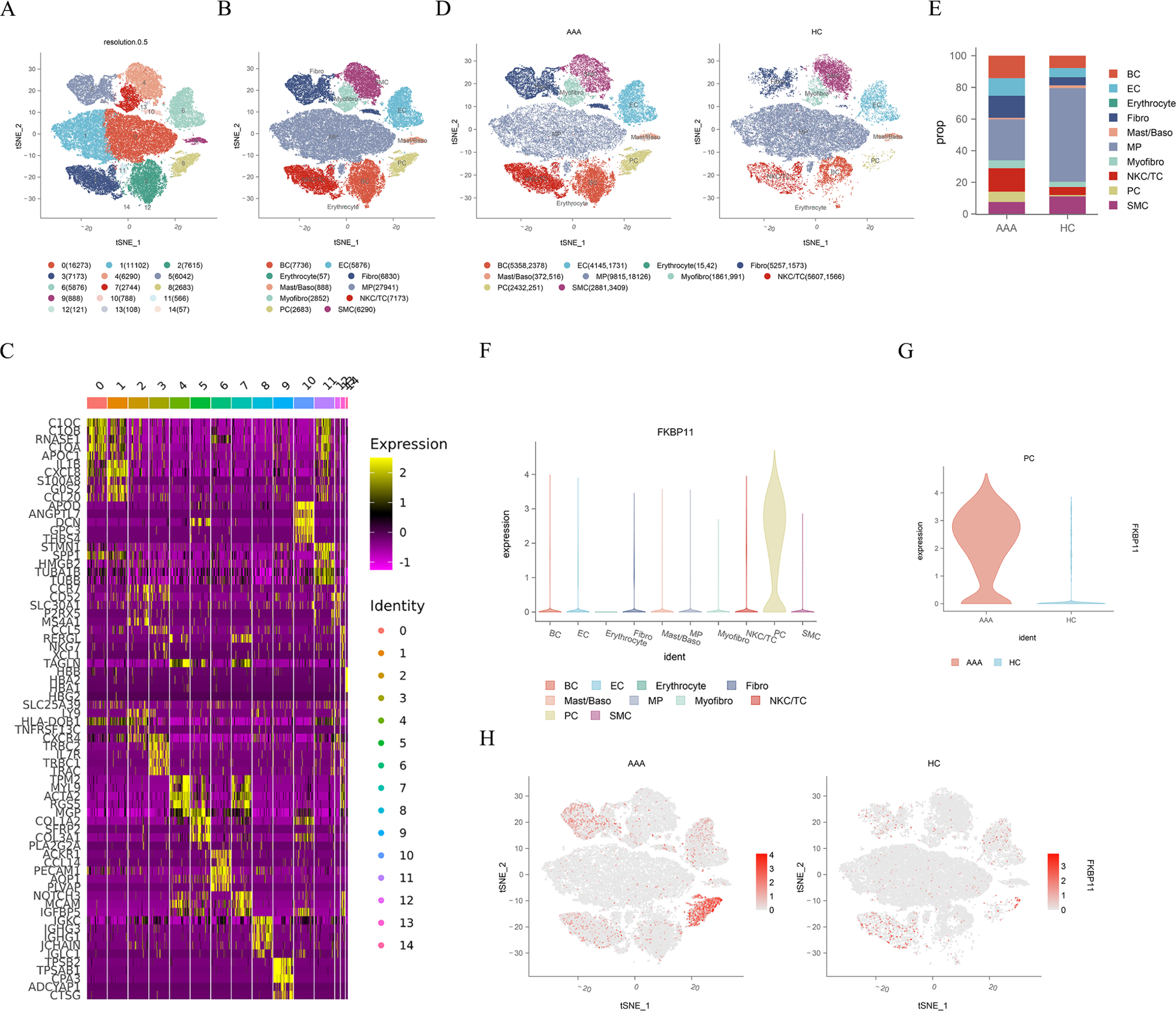
Identification of cell clusters present in the human AAA tissues and healthy aortic samples by ScRNA-seq. (**A**) The t-Distributed Stochastic Neighbor Embedding (t-SNE) plot of all cell clusters at a resolution of 0.5 presented in all samples. (**B**) The t-SNE plot of all cell clusters presented in all samples colored according to identified clusters. (**C**) Heatmap of the top 5 enriched genes in each population in the AAA group.(**D**) The t-SNE plot of cell clusters presented in AAA tissue samples and healthy aortic samples colored according to identified clusters, respectively. (**E**) Percentages of cell populations in the AAA group and the control group. (**F**) The Violinplot of the expression of FKBP11 in different cell clusters. (**G**) The Violinplot of the expression of FKBP11 in PCs from AAA tissue samples and healthy aortic samples. (**H**) The distribution of the expression of FKBP11 in the AAA samples and the control samples.

**Table 5.** The cell types and cell markers detected in AAA tissues and paired peripheral blood samples.

| No | Populations | Cell Markers | Sub-populations |  |
| --- | --- | --- | --- | --- |
| 1 | Mononuclear phagocytes | *LYZ, CD14, CD68, CD163* | Classical monocytes | *LYZ, CD14, FCN1, VCAN, FCGR3A* |
|  |  |  | Non-classical monocytes | *LYZ, FCGR3A, IFITM3* |
|  |  |  | Macrophages | *LYZ, MRC1, CD68, CD163, C1QC* |
|  |  |  | Conventional dendritic cells | *CD1C, CD1E, FCER1A, XCR1* |
| 2 | Endothelial cells | *CDH5, PECAM1, VWF, CLDN5* |  |  |
| 3 | Fibroblasts | *DCN, LUM, COL1A2, COL1A1* |  |  |
| 4 | Mural cells | *ACTA2, TAGLN, MYLK, MYH11, RGS5, MCAM, PDG, FRB* | Myofibroblasts | *RGS5,MCAM,PDGFRB,NOTCH3* |
|  |  |  | Smooth muscle cells | *ACTA2,TAGLN,MYLK,MYH11* |
| 5 | B cells | *MS4A1, CD79A, CD79B* | Naive B cells | *TCL1A, FCER2, IGHM, IGHD* |
|  |  |  | Memory B cells | *CD27, SPIB, AIM2, IGHG1* |
|  |  |  | Germinal center B cells | *NEIL1, AICDA, RGS13, GCSAM* |
| 6 | Plasma cells | *CD79A, JCHAIN, MZB1, IGHG1* | IGHG1-Plasma cells | *CD79A, JCHAIN, MZB1, IGHG1* |
|  |  |  | IGHA1-Plasma cells | *CD79A, JCHAIN, MZB1, IGHA1* |
| 7 | Natural killer cells | *CD3D-, KLRD1, NKG7, KLRC1, FCGR3A* |  |  |
| 8 | T cells | *CD2, CD3D/E/G, TRAC, TRBC1* | T helper cells | *CD4, TNF, IFNG, CCL4, ZFP36L2, GATA3, TNFRSF4, CD40LG* |
|  |  |  | CD8+ effector T cell | *CD3D, CD8A/B, NKG7, GZMA, GNLY* |
|  |  |  | CD4+ naive T cells | *CD4, CCR7, LEF1, SELL, TCF7* |
|  |  |  | CD8+ naive T cells | *CD8A, CCR7, LEF1, SELL, TCF7* |
|  |  |  | Regulatory T cells | *CD4, FOXP3, IL2RA, CTLA4, IKZF2* |
| 9 | Basophils | *LYZ, CLC, GATA2, CPA3, MS4A2* |  |  |
| 10 | Mast cells | *TPSAB1, TPSB2, CPA3* |  |  |
| 11 | Erythrocytes | *HBB, HBA2, HBA1, HBG2* |  |  |

Subsequently, we analyzed the distribution of cell populations in AAA tissue samples and healthy aortic samples **(Figure 7D** and **7E**). In both AAA tissue samples and the healthy control group, mononuclear phagocytes constituted the main immune cell population. However, the proportion of other immune cell populations, such as natural killer cells/T cells (NKC/TC), B cells, and plasma cells (PCs), showed a significant increase in the AAA groups compared to the healthy controls.

In order to further illustrate the participation of FKBP11 in the progression of AAA, we employed both scRNA-seq and IHC staining techniques for AAA tissues and the healthy samples. First, our scRNA-seq analysis revealed that FKBP11 exhibited predominant expression in PCs among all the cell populations examined.(**Figure 7F**) Notably, PCs within AAA tissues displayed a significantly higher level of FKBP11 expression compared to those in healthy aortic tissues, as evidenced by violinplot and featureplot representations.(**Figure 7G** and **7H**)

To validate and complement the scRNA-seq findings, we performed IHC staining on consecutive AAA tissue sections. Consistent with the scRNA-seq results, we observed increased FKBP11 expression in the PCs infiltrating AAA tissues compared to healthy aortic tissues.(**Figure 8**)

**Figure 8.**
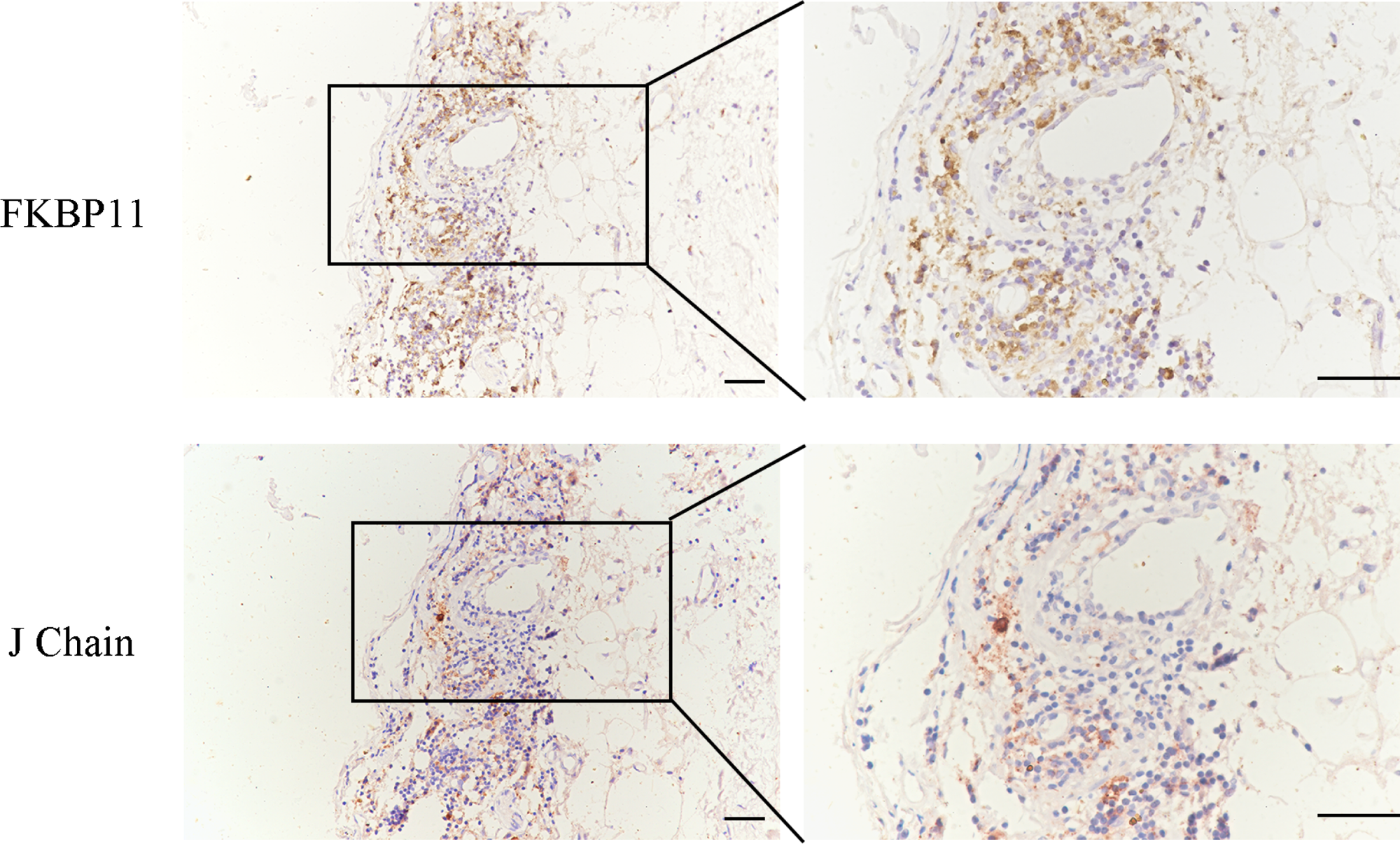
Analysis of the expression of FKBP11 in plasma cells within AAA wall using IHC. The images depict consecutive staining of FKBP11 and J Chain in the representative sections of AAA tissue. Scale bar, 50μm.

### FKBP11(+) Plasma cell exhibited Differential Expression and Functional Enrichment in comparison to FKBP11(-) Plasma Cells

We further sought to delineate the heterogeneity of FKBP11 expression within the plasma cell population. By re-clustering the plasma cells into 8 sub-clusters at a resolution of 0.5, we discerned distinct expression patterns of FKBP11 using featureplot analysis. (**Figure 9A**)

**Figure 9.**
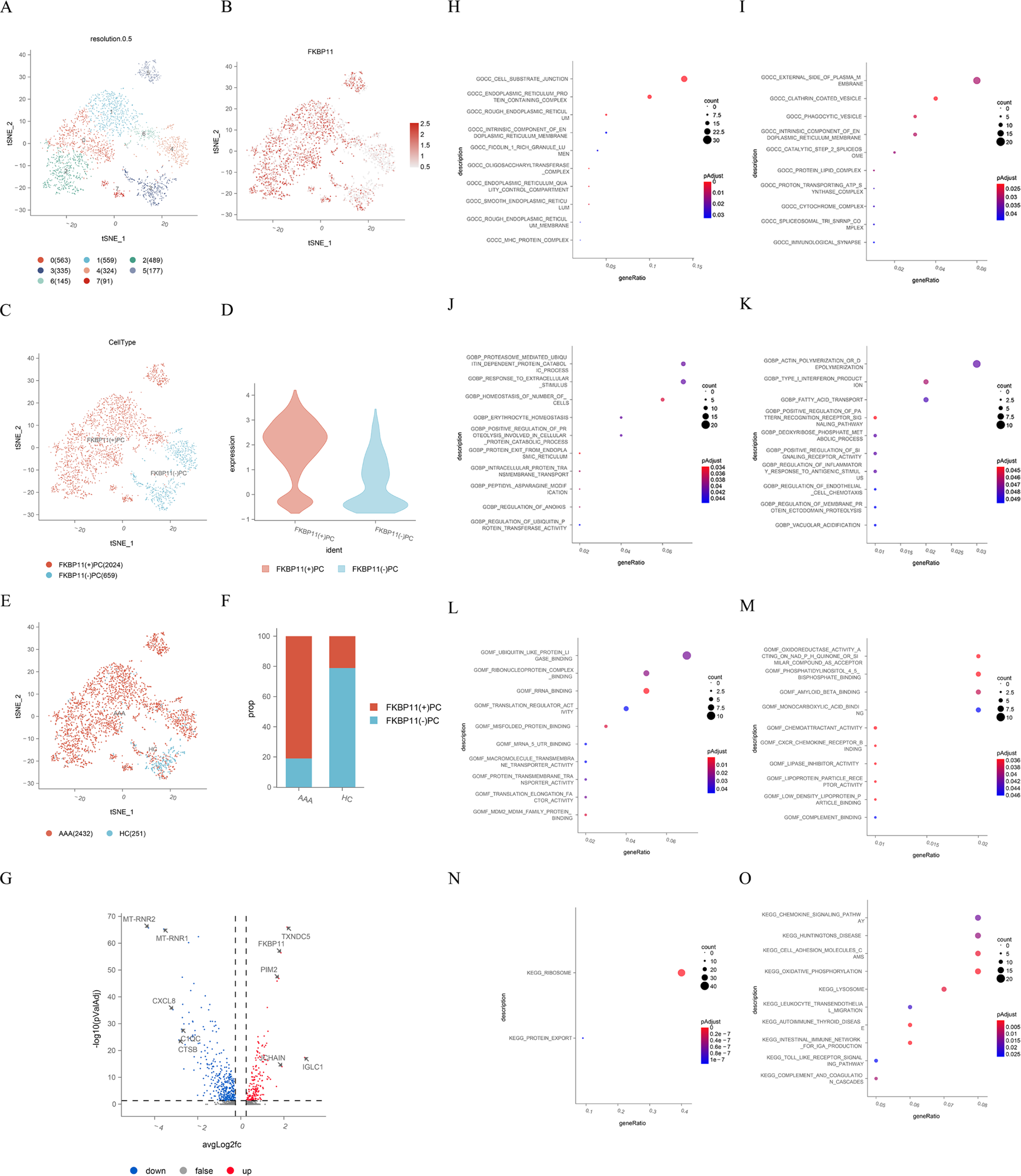
The DEG analysis and functional enrichment analysis on FKBP11(+) PCs and FKBP11(-) PCs. (**A**) The t-SNE plot of sub-populations of PCs colored according to identified sub-clusters. (**B**) The distribution of FKBP11 expression in all sub-clusters as shown in featureplot. (**C**) The t-SNE plot showed the distribution of FKBP11(+)PC and FKBP11(-)PCs. (**D**) Violin plot exhibited the expression levels of FKBP11 in FKBP11(+)PCs and FKBP11(-) PCs. (**E**) The t-SNE plot showed the distribution of PCs in the AAA group and control group. (**F**) Percentages of FKBP11(+)PCs and FKBP11(-) PCs in the AAA group and the control group, respectively. (**G**) The volcano map visually DEGs, with blue and red colors denoting genes expressed at low and high levels, in FKBP11(+) PCs compared with FKBP11(-) PCs. (**H**) and (**I**) GO analysis of the up-regulated and down-regulated functional enrichment in therm of cellular components in FKBP11(+) PCs compared with FKBP11(-) shown in the dotplot. (**J**) and (**K**) GO analysis of the up-regulated and down-regulated functional enrichment in therm of biological processes in FKBP11(+) PCs compared with FKBP11(-) shown in the dotplot. (**L**) and (**M**) GO analysis of the up-regulated and down-regulated functional enrichment in therm of molecular functions in FKBP11(+) PCs compared with FKBP11(-) shown in the dotplot. (**N**) and (**O**) KEGG analysis of the up-regulated and down-regulated enriched signal pathways of FKBP11(+) PCs compared with FKBP11(-) shown in the dotplot.

Specifically, we identified six sub-clusters (Cluster 0, Cluster 1, Cluster 2, Cluster 5, Cluster 6, and Cluster 7) characterized by higher levels of FKBP11 expression, designating them as FKBP11 positive (+) plasma cells. Conversely, two sub-clusters (Cluster 3 and Cluster 4) exhibited lower levels of FKBP11 expression, classifying them as FKBP11 negative (-) plasma cells. (**Figure 9B-9D**)

In our investigation, we observed a notable increase in the abundance of plasma cells (PCs) within AAA tissue samples compared to the control group. Furthermore, the AAA tissue samples exhibited a significantly higher proportion of FKBP11(+) PCs, as illustrated in **Figure 9E** and **9F**.

Through a rigorous analysis, we identified a total of 554 differentially expressed genes (DEGs) between FKBP11(+) and FKBP11(-) plasma cells. Among these DEGs, 184 genes were upregulated, while 370 genes were downregulated. The volcano map visually represents these DEGs, with blue and red colors denoting genes expressed at low and high levels, respectively, in FKBP11(+) PCs compared with FKBP11(-) PCs. (**Figure 9G**)

We proceeded with GO analysis and KEGG analysis to unravel the functional implications of these DEGs (**Figure 9H-9O**). For FKBP11(+) plasma cells, in terms of cellular components, these DEGs were enriched in ribosomal subunits and endoplasmic reticulum protein-containing complexes. Additionally, the upregulated DEGs were found to be significantly enriched in several biological processes, including cytoplasmic translation, protein localization to the endoplasmic reticulum, response to endoplasmic reticulum stress, and response to unfolded protein. As for molecular functions, all of DEGs were associated with ubiquitin protein transferase regulator activity, structural constituent of ribosomes, and misfolded protein binding. Furthermore, the KEGG pathway analysis revealed significant enrichment in protein export and ribosomal pathways. In summary, the heightened expression of FKBP11 in plasma cells suggests an activation of ribosomal and endoplasmic reticulum functions, which are intricately linked to protein translation and export processes.

### Identification of TF-Target Gene Pairs Using SCENIC Analysis in FKBP11(+) and FKBP11(-) Plasma Cells

To elucidate the transcriptional distinctions between FKBP11(+) and FKBP11(-) plasma cell subclusters, we conducted SCENIC analysis to explore potential regulon activities, including transcription factors (TFs) and their target genes, within these two subclusters.

Based on 325 identified regulon activities, specific regulons in FKBP11(+) and FKBP11(-) plasma cell subclusters exhibited clear differences. The top five specific TFs in FKBP11(+) plasma cells were identified as ATF6B, EOMES, FOXO4, ZBTB11, and ZNF12, while key TFs in FKBP11(-) plasma cells were determined to be IRF8, PPARG, KLF4, NFATC2, and STAT6 (**Figure 10A** and **10B**). **Figure 10C** and **10D** depict the PPI networks constructed using TF-target gene pairs for FKBP11(+) and FKBP11(-) plasma cells, respectively, with the key TFs positioned at the core of the networks.

**Figure 10.**
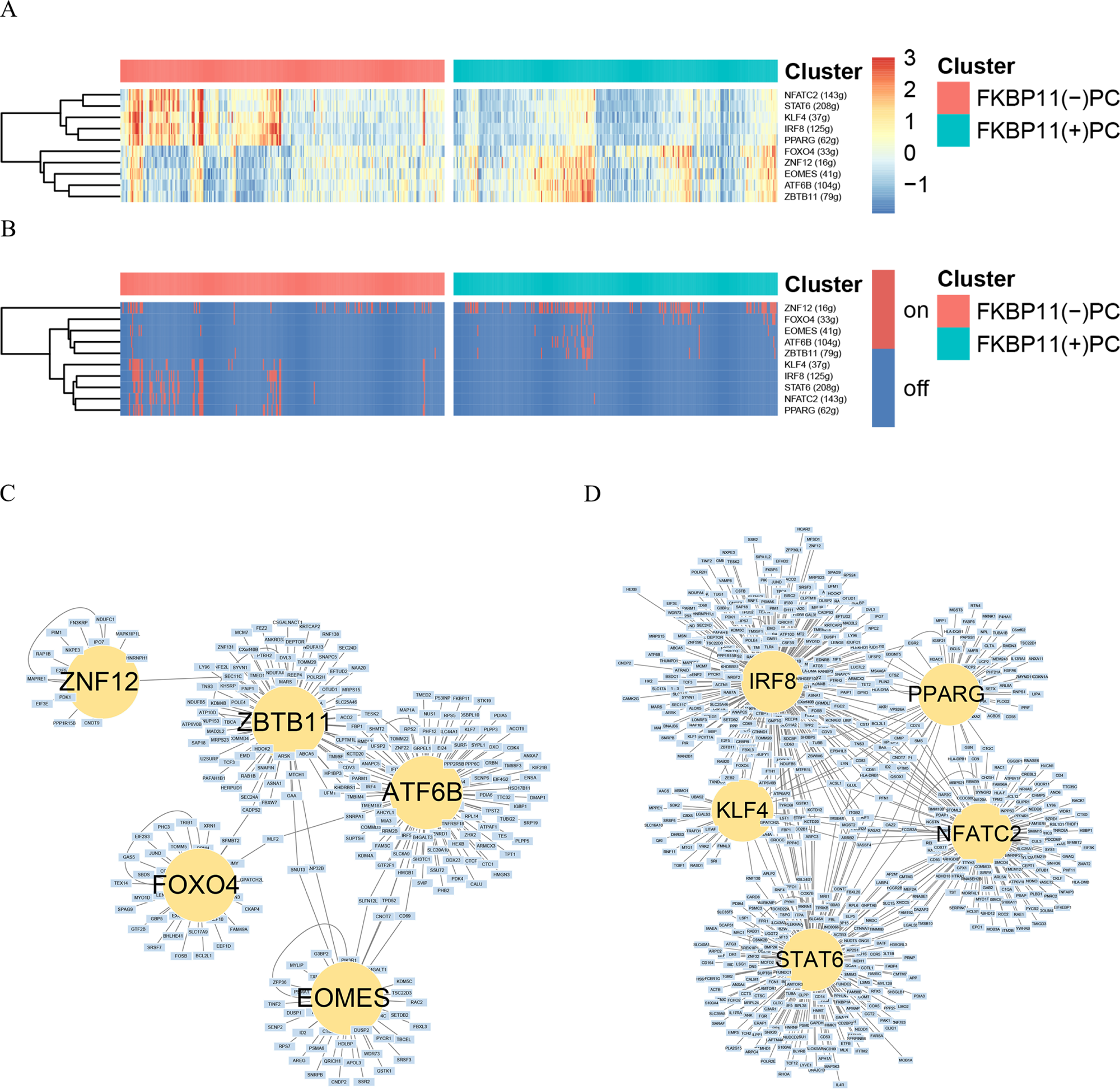
Identification of TF-Target Gene Pairs Using SCENIC Analysis in FKBP11(+) and FKBP11(-) Plasma Cells. (**A**) The heatmap of the top 5 specific regulons in FKBP11(+) PCs and FKBP11(-) PCs based on AUC value. (**B**) The heatmap of the top 5 specific regulons in FKBP11(+) PCs and FKBP11(-) PCs based on the on/off of regulon activity. (**C**) Construction of PPI network of Top 5 regulons and their target genes in FKBP11(+) PCs. (**D**) Construction of PPI network of Top 5 regulons and their target genes in FKBP11(-) PCs.

Notably, our analysis highlighted ATF6B as a key regulator of the high expression of FKBP11 in plasma cells. ATF6B has been associated with endoplasmic reticulum function [10]. Combining this finding with the results of the differential expression analysis, we identified 37 target genes that were up-regulated in FKBP11(+) plasma cells, with a significant enrichment in intrinsic apoptotic signaling and protein processing in the endoplasmic reticulum (**Figure 11A** and **11B**). Conversely, in FKBP11(-) plasma cells, 126 target genes exhibited significant up-regulation. GO and KEGG analyses revealed that these target genes were mainly related to antigen processing and presentation via MHC class II, according with the function of plasma cells (**Figure 11C** and **11D**).

**Figure 11.**
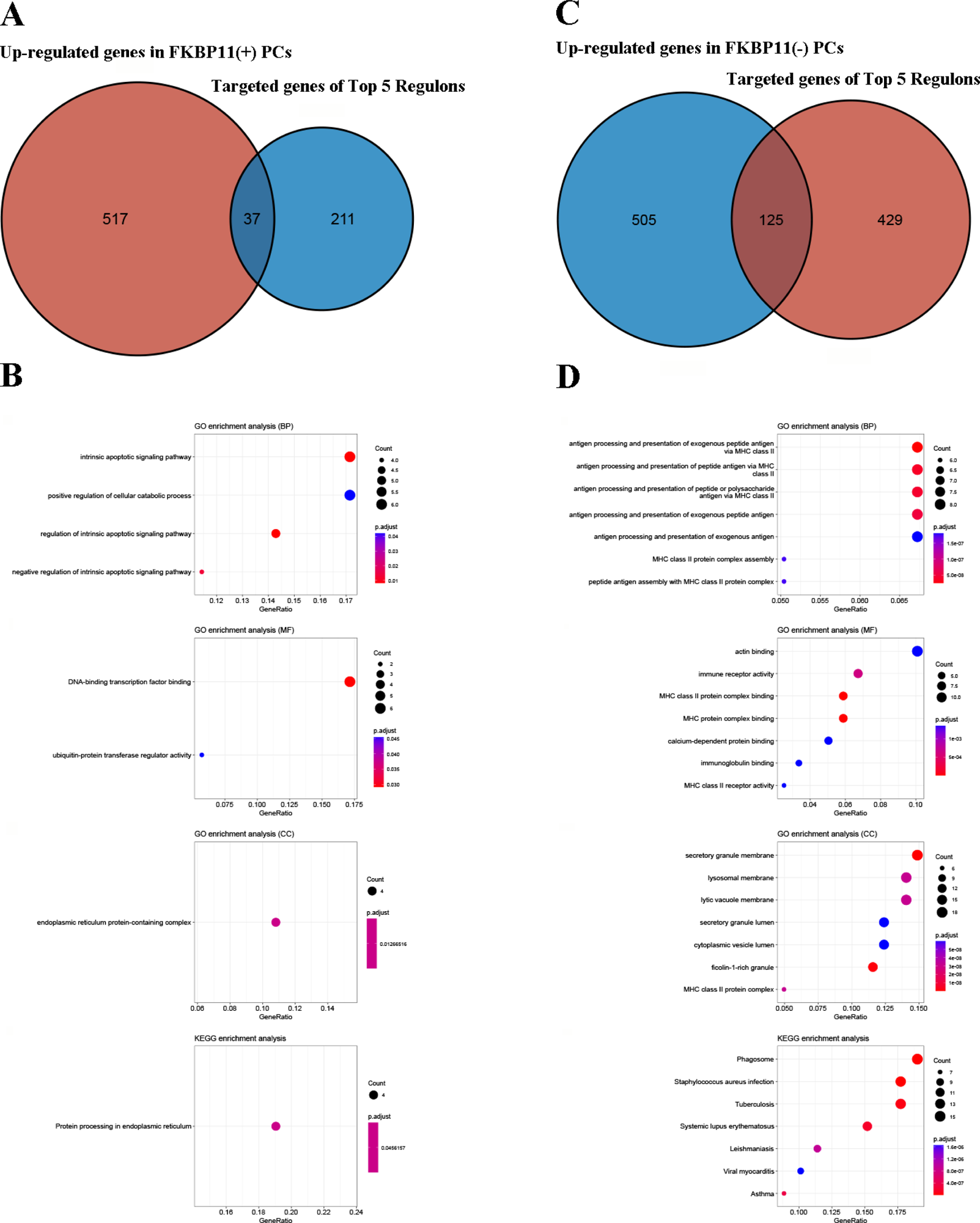
The major functions of TF-Target genes in FKBP11(+) and FKBP11(-) Plasma Cells. (**A**) The Venn diagram of up-regulated DEGs and TF-target genes in FKBP11(+) PCs. (**B**) The GO analysis and KEGG analysis of up-regulated TF-target genes in FKBP11(+) PCs. (**C**) The Venn diagram of up-regulated DEGs and TF-target genes in FKBP11(-) PCs. (**D**) The GO analysis and KEGG analysis of up-regulated TF-target genes in FKBP11(-) PCs.

### Pseudotime Analysis Reveals Phenotypic Switch of Plasma Cells from FKBP11(-) to FKBP11(+) in AAA Vessel Walls

To gain deeper insights into the differentiation process of plasma cells infiltrating AAA tissues, we employed “Monocle 3” R package to calculate trajectories of these cells. (**Figure 12 A-12C**) FKBP11(-) plasma cells were positioned at the initial stage of the trajectory curve, while FKBP11(+) plasma cells were predominantly localized at the terminal stage. To examine the changes in gene expression from the initial cell state (FKBP11(-)PCs) to the terminal state (FKBP11(+) PCs) during differentiation and development, we identified a total of 9 gene expression modules based on the differential expression analysis of pseudotime. (**Figure 12D** and **12E**)

**Figure 12.**
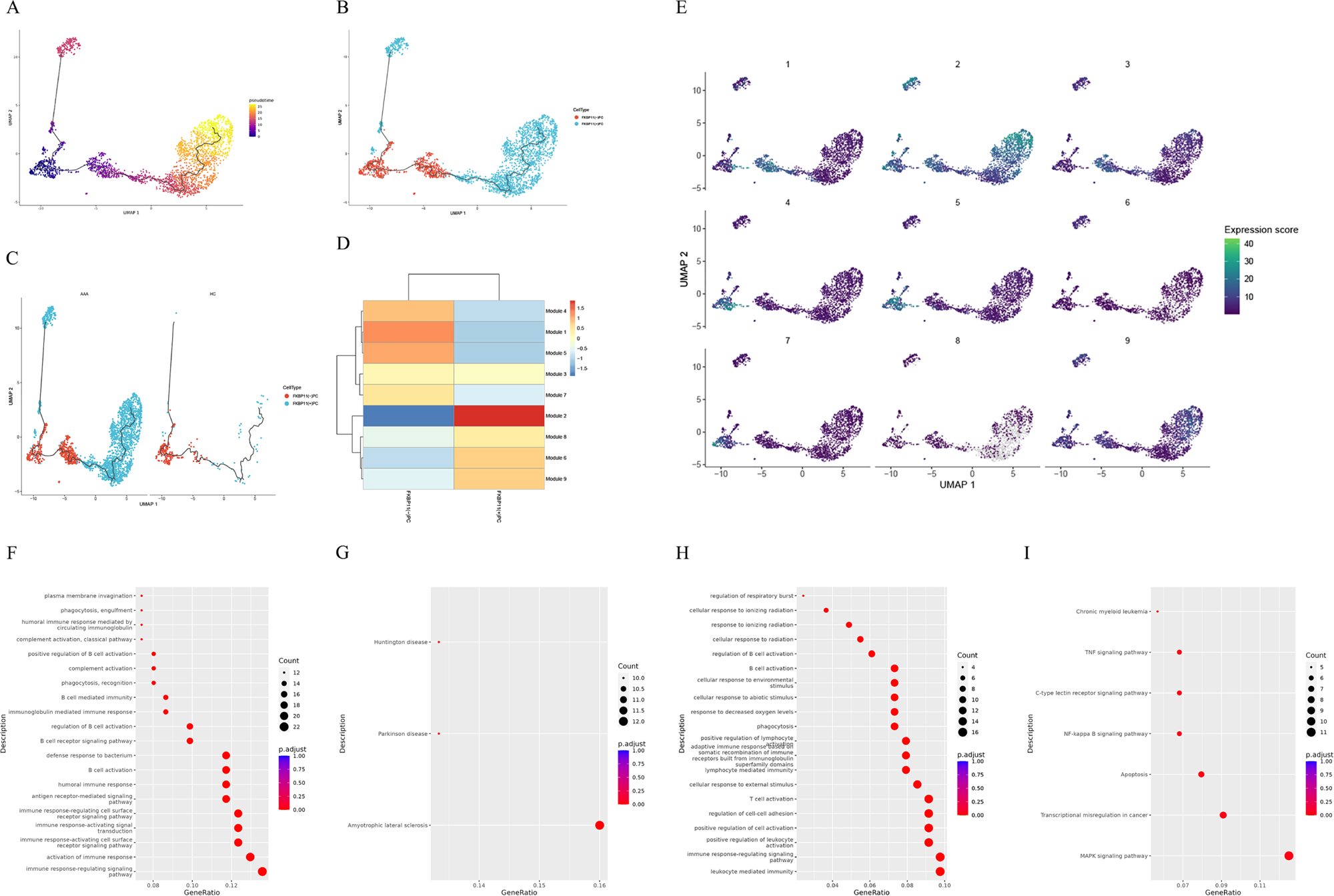
Pseudotime Analysis of Plasma Cells in AAA Vessel Walls. (**A**) The single-cell trajectory of plasma cells predicted by Monocle 3 and ordered in pseudotime. (**B**) The distribution of FKBP11(+) and FKBP11(-) plasma cells in the pseudotime trajector colored according to identified clusters. (**C**) The distribution of FKBP11(+) and FKBP11(-) plasma cells in the pseudotime trajectory split by groups. (**D**) The heatmap plot of gene modules in the two celltypes. (**E**) The single-cell trajectory of plasma cells split by different gene modules. Major GO enrichments (**F**) and KEGG enrichments (**G**) of genes from Module 1. Major GO enrichments (**H**) and KEGG enrichments (**I**) of genes from Module 2.

The subsequent enrichment analysis of these gene modules shed light on the functional and cellular states of plasma cells during different developmental stages. (**Figure 12F-12I**) Specifically, FKBP11(-) plasma cells were enriched in Module 1. GO analysis revealed that these modules were associated with the activation of plasma cells, particularly with a focus on antigen presentation.

In contrast, FKBP11(+) plasma cells were primarily associated with Module 2, which demonstrated enrichments related to T cell activation, leukocyte-mediated immunity, and immune response-regulating signaling pathways.

Based on these findings, we can deduce that a phenotypic switch of plasma cells occurs in the context of AAA, with a transition from FKBP11(-) to FKBP11(+) state. During this differentiation process, the capacity of PCs to mediate lymphocyte immunity becomes enhanced.

### Identification of CD74-APP as the Critical Ligand-Receptor Pair Between FKBP11(+) Plasma Cells and T Cells in AAA Vessel Tissue Samples

The Pseudotime analysis conducted earlier revealed that the phenotypic switch from FKBP11(-) to FKBP11(+) plasma cells in AAA vessel walls correlated with an enhanced ability to mediate lymphocyte immunity. Building upon these findings, we further investigated the explicit interactions between the two sub-populations of PCs and T cells within the AAA microenvironment.

Through re-clustering of the PC population and the NK/T cell population into 8 sub-clusters at a resolution of 0.5, we successfully distinguished distinct sub-populations. These included two PC sub-populations, denoted as Cluster 4 and Cluster 5, which were divided based on the expressions of FKBP11. The remaining clusters were categorized as Naive T cells, Helper T (Th) cells, Regulatory T cells (Treg), Effector T cells (Teff), and NK cells (NKCs). (**Figure 13A** and **13B**)

**Figure 13.**
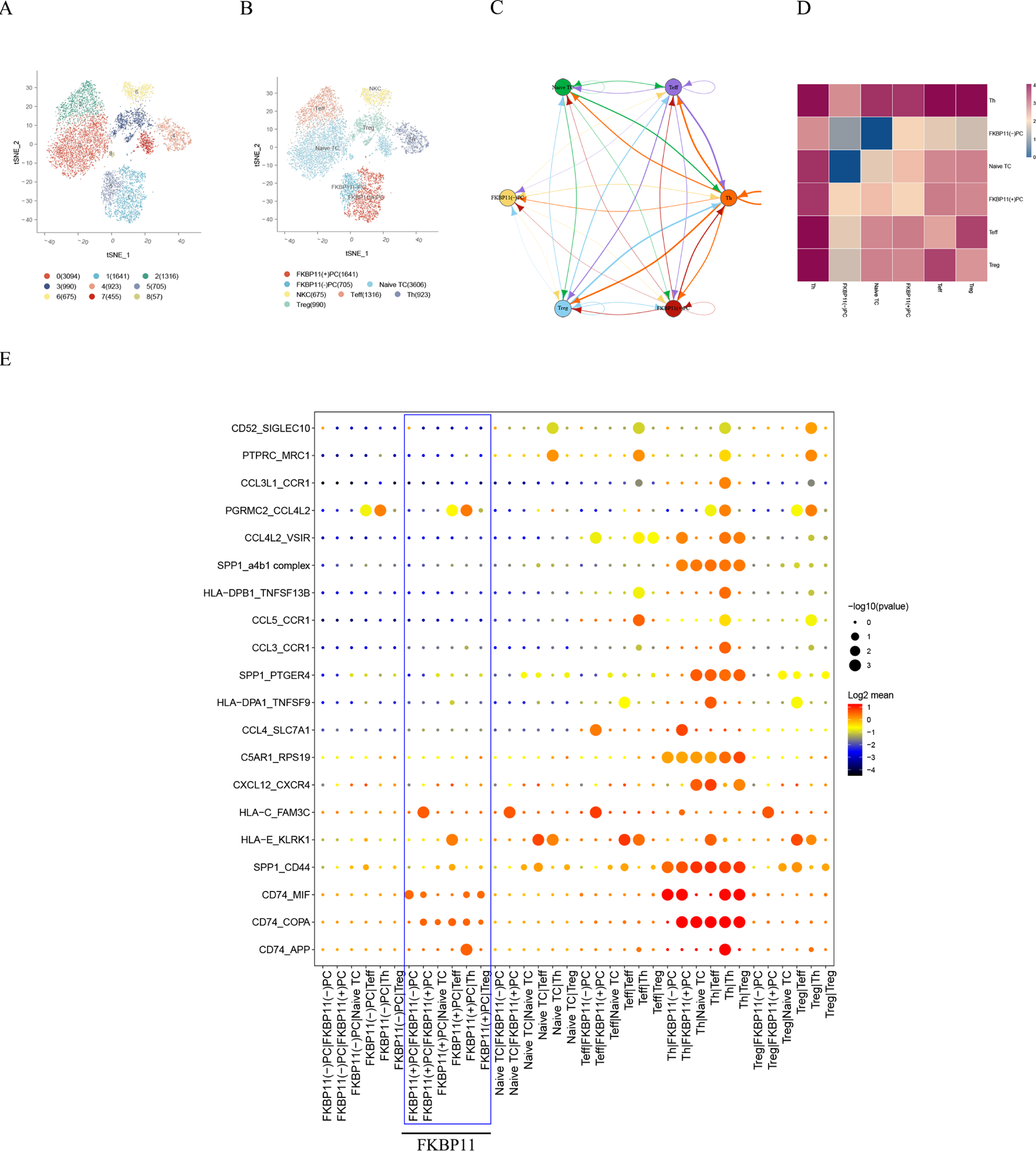
Cell-cell communication among sub-populations of plasma cells and T cells. (**A**) The t-Distributed Stochastic Neighbor Embedding (t-SNE) plot visualized all cell sub-clusters of plasma cells and nature killer cells/T cells at a resolution of 0.5 presented in all samples. (**B**) The t-SNE plot of all cell clusters presented in all samples colored according to identified clusters. (**C**) The interaction network plot and (**D**) the Heatmap plot showed the number and intensity of ligand–receptor pairs among cell sub-types. (**E**) Dotplot of the predicted interactions between plasma cell sub-types and T cell sub-types.

To investigate the cell-cell interaction network among these TC and PC sub-populations, we utilized CellPhoneDB. Notably, we observed an increased number of connections between FKBP11(+) PCs and the various sub-populations of TCs, as compared to FKBP11(-) PCs. In particular, FKBP11(+) PCs exhibited notably stronger interactions with Th cells when compared to other sub-populations of T cells.(**Figure 13C** and **13D**)

The subsequent bubble plots illustrated the Top 20 specific ligand-receptor pairs among these cell types. Of significant interest, we found that CD74, expressed in FKBP11(+) PCs, primarily interacted with Th cells through APP, COPA, and MIF. (**Figure 13E**)

## Discussion

AAA is a degenerative vascular dilatory disease associated with significant morbidity and mortality rates. The complex cellular components and pathogenesis pose challenges for effective medical treatment to prevent AAA development and progression. M6A modification has been implicated in various biological processes, and the m6A level was up-regulated in AAA tissue samples [8]. However, the role of m6A methylation in AAA progression remains unclear. Therefore, it is necessary to conduct a transcriptome-wide analysis to identify altered RNA m6A profiles in AAA tissue samples.

In this study, we detected m6A modifications in AAA tissue samples and healthy controls. We identified 5947 peaks that were common to both groups, accounting for 71.68% of all peaks, suggesting that most m6A modifications are involved in maintaining cellular metabolism. Furthermore, we performed systematic functional analysis of genes with differential m6A modifications and their associations with AAA pathogenesis using GO analysis and KEGG pathway analysis. We found significant enrichments of up-regulated and down-regulated genes, as well as overlapping biological processes in both the AAA and control groups, such as “Ubiquitin mediated proteolysis,” “TGF-beta signaling pathway,” and “Protein processing in endoplasmic reticulum.” However, the m6A levels of these DMGs were distinct between AAA samples and healthy aortic samples. These findings provide valuable insights into the complex dynamic regulatory network of m6A methylation in AAA.

A significant contribution of our study is the development and validation of a risk signature based on AAA m6A-related mRNA. We conducted integrated bioinformatics analysis using MeRIP-seq and existing AAA RNA-seq data from the GEO database to identify hub DMGs and DMRs in AAA, which formed the basis of AMRMS. Moreover, AMRMS was validated as a reliable and powerful gene set for predicting the risk of AAA dilation. Follow-up data from AAA patients in the UK Multicenter Aneurysm Screening Study (MASS) revealed that AAA with a diameter >5.5 cm was associated with a high incidence of rupture [11]. According to the guideline, a maximum diameter exceeding 5.5 cm is considered a critical surgical indication for AAA [2]. However, there is currently no effective targeted drug to inhibit AAA progression in patients whose AAA diameter is below the surgical threshold. ROC curve analysis confirmed that AMRMS was a candidate risk factor associated with large AAA in GSE57691 and GSE98278. Our research enhances the current understanding of AAA and offers a potential tool for risk prediction and personalized management. As a result, it may pave the way for the development of targeted therapies that could effectively inhibit AAA progression and improve patient outcomes.

Among the genes in AMRMS, FKBP11 showed the highest regression coefficient and was considered the most important gene associated with AAA progression. Therefore, we further investigated the potential role of FKBP11 in AAA using scRNA-seq, which provides an unbiased characterization of the in vivo molecular profile and heterogeneity of the complex cellular components in the AAA vessel wall. Our scRNA-seq analysis revealed a significant up-regulation of FKBP11 mRNA in AAA tissue samples, predominantly in plasma cells. This finding provided new insights into the function of plasma cells in AAA.

Accumulating evidence suggests that human AAA exhibits significant autoimmune features, including the secretion of autoantibodies and dysfunction of regulatory T cells [12–15]. Previous studies have reported an increase in B cell/plasma cell populations in human AAA, and the depletion of B cells has been shown to be protective [13, 16, 17]. In autoimmune diseases, plasma cells differentiate from B cells and serve as major producers of autoantibodies. FKBP11 is a plasma cell-specific antibody folding catalyst that is up-regulated in response to endoplasmic reticulum (ER) stress [18, 19]. It regulates the unfolded protein response (UPR) to accommodate the increased burden of nascent immunoglobulins and prevent the accumulation of misfolded proteins in the ER [20, 21]. In our research, pseudotime analysis indicated a phenotypic switch from FKBP11(-) to FKBP11(+) resident plasma cells in AAA walls, while plasma cells in healthy aortas from donors predominantly exhibited the FKBP11(-) phenotype. These findings suggest that not only an increased number of plasma cells, but also the activation of ER-related functions in plasma cells, play important roles in AAA pathogenesis.

Another noteworthy finding from the pseudotime analysis was that with increased expression of FKBP11, plasma cells exhibited enrichment in functions related to mediating T cell activation, as evidenced by changes in gene modules. Furthermore, cell-cell communication analysis revealed increased interactions between FKBP11(+) plasma cells and T cell subpopulations compared to FKBP11(-) plasma cells. We identified several highly enriched receptor-ligand pairs, including CD74-COPA/MIF/APP, involved in the interaction between T cells and plasma cells. The exact role of these receptor-ligand pairs in T cell-plasma cell interactions remains unclear. CD74, also known as the HLA-DR antigens-associated invariant chain, acts primarily as a chaperone for MHC class II molecules and is highly expressed in immune cells. It also participates in the transport of other non-MHC-II proteins [22]. CD74, an integral membrane protein, is associated with the promotion of B cell growth and survival [23]. Milatuzumab, a humanized anti-CD74 monoclonal antibody, has been shown to decrease CD20/CD74 aggregates and cell adhesion, leading to cell death [24, 25]. Our findings suggest that the drug targeting on CD74, like Milatuzumab, may potentially prevent AAA progression by suppressing the activation of FKBP11(+) plasma cells.

It is important to acknowledge the limitations of our study. While we provide evidence for the involvement of m6A methylation in AAA, further investigations are needed to elucidate the specific mechanisms through which m6A methylation affects the identified DMGs and contributes to AAA pathogenesis. The rarity of clinical specimens of human AAA makes it challenging to extract and culture a sufficient number of primary resident plasma cells to identify potential mechanisms involved in AAA pathogenesis.

### Conclusions

Our study highlights the importance of m6A methylation in AAA pathogenesis and presents a comprehensive transcriptome-wide map of m6A modifications in AAA tissues. Using various bioinformatics and machine learning algorithms, we developed a novel and powerful m6A-related signature, AMRMS, for predicting AAA progression. Furthermore, through scRNA-seq, we discovered that FKBP11, the most specific risk factor in AMRMS, is primarily expressed in plasma cells within AAA walls and is involved in ER stress. Our research sheds light on the role of plasma cells in AAA and suggests that they may be potential targets for treating AAA. Our findings may provide valuable insights for diagnosing and treating AAA, with the aim of preventing AAA dilation in patients in the future.

## Acknowledgements

We thank Mitchell Arico from Liwen Bianji (Edanz) (https://www.liwenbianji.cn) for editing the language of a draft of this manuscript.

## Authors’ contributions

YCH, and JX performed experiments and data analysis. YCH and JZ wrote the article. SJX, and YL provided technical support. YSH, PE, SYW, and HJ contributed to the discussion of the project and article. JZ, BD and PE did critical editing. JZ and YCH designed research and discussed results.

## Abbreviations

AAA: abdominal aortic aneurysm
AMRMS: AAA m6A-related mRNA signature
AUC: Area Under the Curve
BP: biological process
CBBs: Cell Barcoded Magnetic Beads
CC: cellular component
CCA: Canonical Correlation Analysis
CMUaB: CMU Aneurysm Biobank
CDS: coding sequence
CTA: computed tomography angiography
DEGs: differential expressed genes
DMGs: differentially methylated genes
ECs: endothelial cells
Enet: Elastic Network
ER: endoplasmic reticulum
FC: Fold Change
GBM: Generalized Boosted Regression Modeling
glmBoost: Generalized Linear Model Boosting
GO: gene ontology
HC: healthy control
IHC: Immunohistochemistry
IP: immunoprecipitation
KEGG: Kyoto encyclopedia of genes and genomes
LDA: Linear Discriminant Analysis
LOOCV: Leave-One-Out Cross-Validation
m6A: N6-methyladenosine
MASS: Multicenter Aneurysm Screening Study
MeRIP-seq: methylated RNA immunoprecipitation with next-generation sequencing
MF: molecular function
MPs: mononuclear phagocytes
NKCs/TCs: natural killer cells/T cells
PCs: plasma cells
PCA: principal Component Analysis
PPI: Protein-Protein Interactions
RF: Random Forest
ROC: Receiver Operating Characteristic
SMCs: smooth muscle cells
SMGs: specific methylated genes
Stepglm: Step Generalized Linear Model
SVM: Support Vector Machine
t-SNE: t-Distributed Stochastic Neighbor Embedding
Teff: Effector T cells
Th: Helper T cells
Treg: Regulatory T cells
UMI: unique molecular identifiers
UPR: unfolded protein response
XGBoost: eXtreme Gradient Boosting
3’UTR: 3’ untranslated region
5’UTR: 5’ untranslated region.

## References

1. Sakalihasan N, Limet R, Defawe OD. Abdominal aortic aneurysm. Lancet. 2005;365(9470):1577–1589.

2. Isselbacher EM, Preventza O, Hamilton Black J 3rd, et al. 2022 ACC/AHA Guideline for the Diagnosis and Management of Aortic Disease: A Report of the American Heart Association/American College of Cardiology Joint Committee on Clinical Practice Guidelines. Circulation. 2022;146(24):e334–e482.

3. Jiang X, Liu B, Nie Z, et al. The role of m6A modification in the biological functions and diseases. Signal Transduct Target Ther. 2021;6(1):74.

4. Qin Y, Li L, Luo E, et al. Role of m6A RNA methylation in cardiovascular disease (Review). Int J Mol Med. 2020;46(6):1958–1972.

5. Golledge J. Abdominal aortic aneurysm: update on pathogenesis and medical treatments. Nat Rev Cardiol. 2019;16(4):225–242.

6. Davis FM, Rateri DL, Daugherty A. Abdominal aortic aneurysm: novel mechanisms and therapies. Curr Opin Cardiol 2015;30:566–73.

7. Deng LJ, Deng WQ, Fan SR, et al. m6A modification: recent advances, anticancer targeted drug discovery and beyond. Mol Cancer. 2022;21(1):52.

8. He Y, Xing J, Wang S, Xin S, Han Y, Zhang J. Increased m6A methylation level is associated with the progression of human abdominal aortic aneurysm. Ann Transl Med. 2019;7(24):797.

9. Fu C, Feng L, Zhang J, Sun D. Bioinformatic analyses of the role of m6A RNA methylation regulators in abdominal aortic aneurysm. Ann Transl Med. 2022;10(10):547.

10. Saghaleyni R, Malm M, Moruzzi N, Zrimec J, Razavi R, Wistbacka N, Thorell H, Pintar A, Hober A, Edfors F, Chotteau V, Berggren PO, Grassi L, Zelezniak A, Svensson T, Hatton D, Nielsen J, Robinson JL, Rockberg J. Enhanced metabolism and negative regulation of ER stress support higher erythropoietin production in HEK293 cells. Cell Rep. 2022;39(11):110936.

11. Ashton HA, Buxton MJ, Day NE, Kim LG, Marteau TM, Scott RA, Thompson SG, Walker NM. The Multicentre Aneurysm Screening Study (MASS) into the effect of abdominal aortic aneurysm screening on mortality in men: a randomised controlled trial. Lancet. 2002;360(9345):1531–1539.

12. Jagadesham VP, Scott DJ, Carding SR. Abdominal aortic aneurysms: an autoimmune disease?. Trends Mol Med. 2008;14(12):522–529.

13. Lu S, White JV, Nwaneshiudu I, Nwaneshiudu A, Monos DS, Solomides CC, Oleszak EL, Platsoucas CD. Human abdominal aortic aneurysm (AAA): Evidence for an autoimmune antigen-driven disease. Autoimmun Rev. 2022;21(10):103164.

14. Meng X, Yang J, Dong M, Zhang K, Tu E, Gao Q, Chen W, Zhang C, Zhang Y. Regulatory T cells in cardiovascular diseases. Nat Rev Cardiol. 2016;13(3):167–179.

15. Suh MK, Batra R, Carson JS, Xiong W, Dale MA, Meisinger T, Killen C, Mitchell J, Baxter BT. Ex vivo expansion of regulatory T cells from abdominal aortic aneurysm patients inhibits aneurysm in humanized murine model. J Vasc Surg. 2020;72(3):1087–1096.e1.

16. Schaheen B, Downs EA, Serbulea V, Almenara CC, Spinosa M, Su G, Zhao Y, Srikakulapu P, Butts C, McNamara CA, Leitinger N, Upchurch GR Jr, Meher AK, Ailawadi G. B-Cell Depletion Promotes Aortic Infiltration of Immunosuppressive Cells and Is Protective of Experimental Aortic Aneurysm. Arterioscler Thromb Vasc Biol. 2016;36(11):2191–2202.

17. Spinosa MD, Montgomery WG, Lempicki M, Srikakulapu P, Johnsrude MJ, McNamara CA, Upchurch GR Jr, Ailawadi G, Leitinger N, Meher AK. B Cell-Activating Factor Antagonism Attenuates the Growth of Experimental Abdominal Aortic Aneurysm. Am J Pathol. 2021;191(12):2231–2244.

18. Preisendörfer S, Ishikawa Y, Hennen E, Winklmeier S, Schupp JC, Knüppel L, Fernandez IE, Binzenhöfer L, Flatley A, Juan-Guardela BM, Ruppert C, Guenther A, Frankenberger M, Hatz RA, Kneidinger N, Behr J, Feederle R, Schepers A, Hilgendorff A, Kaminski N, Meinl E, Bächinger HP, Eickelberg O, Staab-Weijnitz CA. FK506-Binding Protein 11 Is a Novel Plasma Cell-Specific Antibody Folding Catalyst with Increased Expression in Idiopathic Pulmonary Fibrosis. Cells. 2022;11(8):1341.

19. Ruer-Laventie J, Simoni L, Schickel JN, Soley A, Duval M, Knapp AM, Marcellin L, Lamon D, Korganow AS, Martin T, Pasquali JL, Soulas-Sprauel P. Overexpression of Fkbp11, a feature of lupus B cells, leads to B cell tolerance breakdown and initiates plasma cell differentiation. Immun Inflamm Dis. 2015;3(3):265–279.

20. Herrema H, Guan D, Choi JW, Feng X, Salazar Hernandez MA, Faruk F, Auen T, Boudett E, Tao R, Chun H, Ozcan U. FKBP11 rewires UPR signaling to promote glucose homeostasis in type 2 diabetes and obesity. Cell Metab. 2022;34(7):1004–1022.e8.

21. Wang X, Cui X, Zhu C, Li M, Zhao J, Shen Z, Shan X, Wang L, Wu H, Shen Y, Ni Y, Zhang D, Zhou G. FKBP11 protects intestinal epithelial cells against inflammation-induced apoptosis via the JNK-caspase pathway in Crohn’s disease. Mol Med Rep. 2018;18(5):4428–4438.

22. Su H, Na N, Zhang X, Zhao Y. The biological function and significance of CD74 in immune diseases. Inflamm Res. 2017;66(3):209–216.

23. David K, Friedlander G, Pellegrino B, Radomir L, Lewinsky H, Leng L, Bucala R, Becker-Herman S, Shachar I. CD74 as a regulator of transcription in normal B cells. Cell Rep. 2022;41(5):111572.

24. Xu S, Zhang X, Chen Y, Ma Y, Deng J, Gao X, Guan S, Pan F. Anti-CD74 antibodies in spondyloarthritis: A systematic review and meta-analysis. Semin Arthritis Rheum. 2021;51(1):7–14.

25. Alinari L, Yu B, Christian BA, Yan F, Shin J, Lapalombella R, Hertlein E, Lustberg ME, Quinion C, Zhang X, Lozanski G, Muthusamy N, Prætorius-Ibba M, O’Connor OA, Goldenberg DM, Byrd JC, Blum KA, Baiocchi RA. Combination anti-CD74 (milatuzumab) and anti-CD20 (rituximab) monoclonal antibody therapy has in vitro and in vivo activity in mantle cell lymphoma. Blood. 2011;117(17):4530–4541.

